# Direct recruitment of Mis18 to interphase spindle poles promotes CENP-A chromatin assembly

**DOI:** 10.1101/2023.07.03.547485

**Authors:** Nitobe London, Bethan Medina-Pritchard, Christos Spanos, Juri Rappsilber, Jeyaprakash Arulanandam, Robin C. Allshire

## Abstract

CENP-A chromatin specifies mammalian centromere identity, and its chaperone HJURP replenishes CENP-A when recruited by the Mis18 complex (Mis18C) via M18BP/KNL2 to CENP-C at kinetochores during interphase. However, the Mis18C recruitment mechanism remains unresolved in species lacking M18BP1, such as fission yeast. Fission yeast centromeres cluster at G2 spindle pole bodies (SPBs) when CENP-A^Cnp1^ is replenished and where Mis18C also localizes. We show that SPBs play an unexpected role in concentrating Mis18C near centromeres through the recruitment of Mis18 by direct binding to the major SPB <u>LI</u>nker of <u>N</u>ucleoskeleton and <u>C</u>ytoskeleton (LINC) complex component Sad1. Mis18 recruitment by Sad1 is important for CENP-A^Cnp1^ chromatin establishment and acts in parallel with a CENP-C-mediated Mis18C recruitment pathway to maintain centromeric CENP-A^Cnp1^, but is independent of Sad1-mediated centromere clustering. SPBs therefore provide a non-chromosomal scaffold for both Mis18C recruitment and centromere clustering during G2. This centromere-independent Mis18-SPB recruitment provides a mechanism that governs *de novo* CENP-A^Cnp1^ chromatin assembly by the proximity of appropriate sequences to SPBs and highlights how nuclear spatial organization influences centromere identity.

## Introduction

Centromeres are chromosomal sites that mediate accurate chromosome segregation during cell division. How centromeres are specified and maintained at a single location on monocentric chromosomes is an unresolved question. Chromatin containing the centromere-specific histone H3 variant CENP-A (Cnp1 in the fission yeast, *Schizosaccharomyces pombe*) underlies kinetochores at many eukaryotic centromeres including those of human chromosomes. Epigenetic mechanisms maintain centromeres by templating CENP-A deposition at sites where CENP-A chromatin was previously assembled^1,2^. However, *de novo* centromere formation at chromosomal locations lacking CENP-A can also occur^3–7^. Such neocentromere formation may contribute to speciation and to oncogenesis^8–11^. Quality control mechanisms hinder neocentromere formation by promoting removal of CENP-A from non-centromeric locations^12–14^. Multiple mechanisms therefore ensure centromere maintenance at their correct locations to promote cell viability.

Maintenance of centromere location is ultimately determined by CENP-A chromatin assembly factor activity. In many species, including *S. pombe,* centromeres are composed of central core domains of high CENP-A^Cnp1^ density flanked by repetitive DNA sequences that form heterochromatin^9^. Centromeres are frequently formed on AT-rich DNA, but underlying sequence alone is generally insufficient to specify centromere formation except for point centromeres such as those of budding yeast^9^. During replication, CENP-A nucleosomes are distributed to both sister chromatids, halving CENP-A levels at centromeres, with histone H3 nucleosomes initially assembled as placeholders^15^. CENP-A^Cnp1^ nucleosomes replace H3 nucleosomes via the activity of the histone chaperone HJURP (*S. pombe* Scm3) and associated factors during G2 in *S. pombe* and telophase/G1 in human cells^16–20^. Centromere-specific recruitment of HJURP^Scm3^ is mediated by the Mis18 complex (Mis18C), which is composed of Mis18, Mis16, Eic1/Mis19 and Eic2/Mis20 in *S. pombe*^21–24^, and hMis18α, hMis18β, and Mis18BP1/KNL2 (M18BP1) in human cells^25–27^. *S. pombe* Mis18 is homologous to both hMis18α, hMis18β, and *S. pombe* Mis16 is equivalent to human RbAp46/48^21, 25^. M18BP1 binds CENP-A, CENP-I, and CENP-C at mammalian kinetochores and consequently recruits Mis18C to centromeres^28–30^. However, *S. pombe* lacks a M18BP1 ortholog, and although *S. pombe* Mis18 directly interacts with CENP-C^Cnp3^ *in vitro*, mutations in Mis18 that disrupt CENP-C^Cnp3^-Mis18 association only modestly reduce CENP-A^Cnp1^ localization at centromeres^24, 25, 31^. Furthermore, *S. pombe* CENP-C^Cnp3^ is not essential, indicating other Mis18C recruitment mechanisms remain to be discovered^32^.

All centromeres associate with the nuclear periphery in a cluster adjacent to the spindle pole body (SPB) in yeasts, and centromere clustering during interphase is widespread in eukaryotic cells including those of human tissues^10, 33–35^. SPBs are the yeast nuclear mitotic microtubule organizing centers - equivalent to metazoan centrosomes^36^. Centromeres and thus kinetochore proteins localize close to SPBs in G2 *S. pombe* cells. SPB-centromere clustering ensures efficient kinetochore microtubule capture in early mitosis, and also promotes efficient CENP-A^Cnp1^ deposition on centromeric DNA^37–41^. Notably, CENP-A^Cnp1^ incorporation depends on spatial proximity of substrate DNA to SPB-centromere clusters^41^. This is consistent with the concentration of Mis18C and Scm3^HJURP^ with clustered centromeres throughout G2, when new CENP-A^Cnp1^ deposition occurs^18, 21, 22, 24, 42, 43^. Compromised Mis18C function results in loss of both CENP-A^Cnp1^ and HJURP^Scm3^ from *S. pombe* centromeres, but Mis18 remains localized when CENP-A^Cnp1^, HJURP^Scm3^ or CENP-C^Cnp3^ function is disrupted^21, 31, 42, 43^. Mis18C components may therefore localize to SPB-centromere clusters independently of centromere proteins.

It remains unclear how SPB-centromere association is mediated. Most *S. pombe* SPB proteins reside on the cytoplasmic plaque on the outer nuclear membrane during interphase, while only a few reside on the inner nuclear membrane (INM)^36^ (Figure 1A). Known INM-associated proteins at SPBs include gamma-tubulin ring complex (γ-TuRC) components, the INM-associated proteins Lem2 and Nur1/Mug154, the spindle- and kinetochore-linking protein Csi1, and the <u>LI</u>nker of <u>N</u>ucleoskeleton and <u>C</u>ytoskeleton (LINC) complex protein Sad1^38, 44–47^. G2 SPB-centromere clustering is moderately disrupted in cells lacking Csi1 and is exacerbated by loss of Lem2^38, 48, 49^. Sad1 links centromeres to the nuclear periphery, and a proportion of *sad1* mutant cells completely dissociate all centromeres from SPBs^50, 51^. Defective kinetochore function perturbs SPB-centromere association, suggesting that Sad1 or Csi1 may interact directly with kinetochore proteins to mediate SPB-centromere connections^38, 52, 53^. Although SPB-centromere linkages depend on Sad1, the protein interaction network at the INM remains poorly understood.

**Figure 1.**
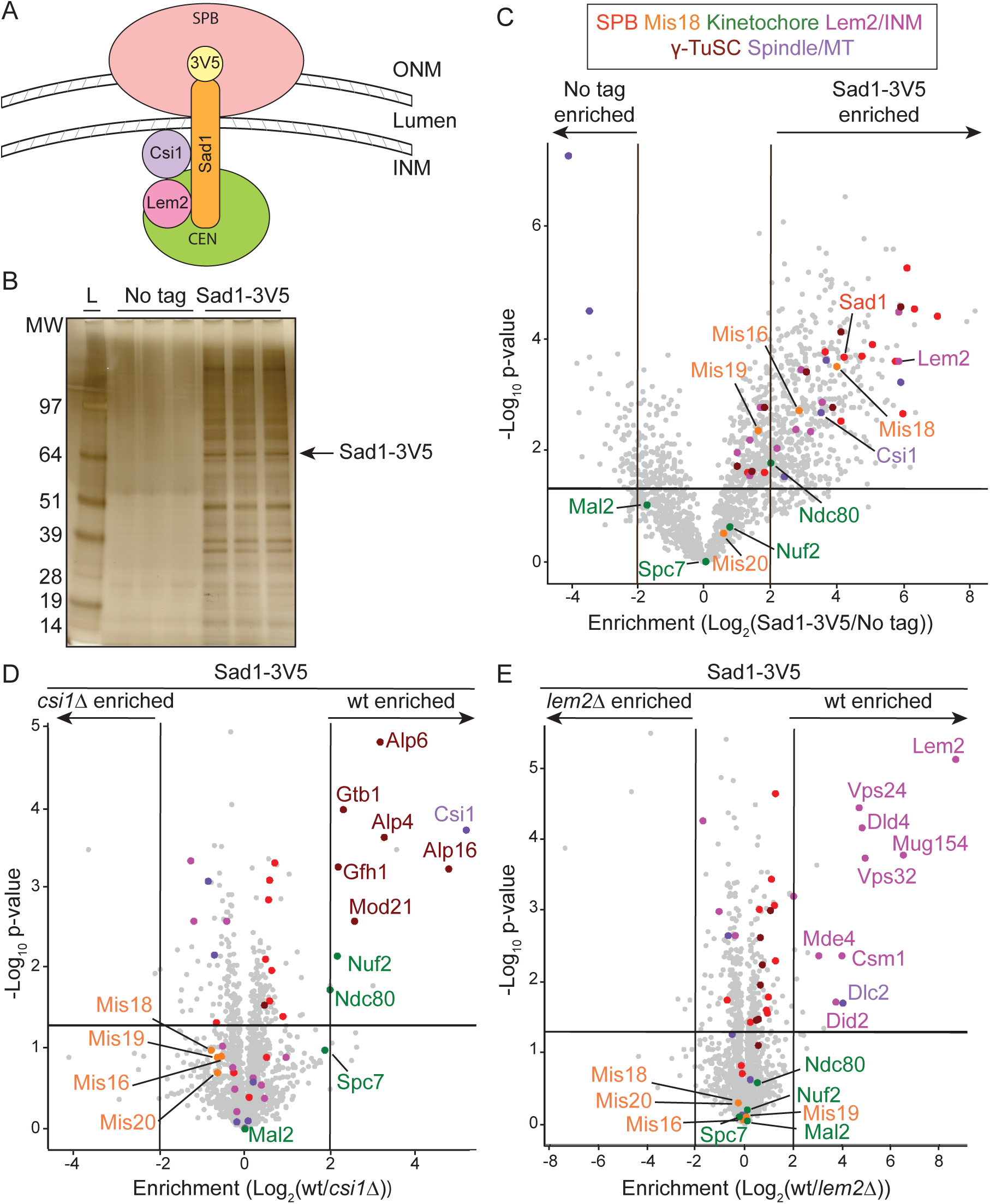
Centromere-, INM-, and spindle-related proteins copurify with Sad1. (A) Diagram of interphase *S. pombe* spindle pole body (SPB) association with the nuclear membrane, showing Sad1 localization at the inner nuclear membrane (INM), its association with centromeres, and its interaction with Csi1 and Lem2. Centromeres of all three chromosomes cluster at SPBs. ONM: Outer Nuclear Membrane. (B) Silver stained SDS-PAGE gel of three independent anti-V5 IPs from extracts of untagged or *sad1-3V5* cells used for LFQ-MS. L: MW ladder, kDa. (C) Volcano plot of LFQ-MS data comparing proteins enriched in anti-V5 Sad1-3V5 IPs relative to untagged control IPs. Lower cutoff corresponds to p = 0.05 and threshold bars are placed at Log_2_(2) (equal to 4-fold enrichment). Samples compared by Student’s t-test with alpha = 0.05 (see Methods). Dots for proteins in select categories color coded as indicated (Table S1). (D, E) Volcano plots of LFQ-MS data comparing proteins enriched with Sad1-3V5 in *csi1*Δ or *lem2*Δ cells relative to wild-type (wt) cells. Dot colors as in (C).

To better characterize *S. pombe* SPB-centromere association we adopted an *in vivo* cleavage approach to identify proteins that specifically associate with the nucleoplasmic region of Sad1. This strategy, along with biochemical and structural analyses, revealed that Mis18 directly interacts with the first 60 residues of Sad1. Our separation-of-function *sad1-4A* mutant causes Mis18, but not centromeres, to dissociate from SPBs. This mutant revealed that recruitment of Mis18 by Sad1 to SPBs promotes *de novo* CENP-A^Cnp1^ chromatin establishment on naïve centromere DNA and contributes to endogenous CENP-A^Cnp1^ chromatin maintenance. Our analyses show that this Mis18-Sad1 recruitment pathway at SPBs also operates in parallel to a centromere-based Mis18 recruitment pathway to maintain existing centromeres. Thus, we uncover a novel mechanism for Mis18C recruitment that does not require direct recruitment of Mis18 to kinetochores for CENP-A deposition at centromeres. Rather, Mis18C operates through a SPB-based platform distinct from kinetochores and even chromosomes, highlighting the importance of the spatial positioning of centromeres within nuclei for centromere identity and integrity.

## Results

### SPB, INM, and kinetochore proteins associate with Sad1

The connection of interphase SPBs with centromeres depends on Sad1, but the full set of proteins that reside on the nucleoplasmic face of SPBs with Sad1 is not known (Figure 1A). To identify Sad1-associated proteins, immunoprecipitated (IP) Sad1-3V5 (C-terminally 3xV5-tagged) from cell extracts was analyzed by label-free quantitative mass spectrometry (IP/LFQ-MS) and compared with untagged control cell IPs (Figures 1B-C). A complete set of structural SPB proteins, components of the INM, γ-TuRC, kinetochore proteins, the Mis18 complex (Mis18C), and spindle-associated proteins were enriched with Sad1-3V5 (Figure 1C; Tables S1 and S2).

Csi1 and Lem2 associate with Sad1 and contribute to centromere clustering at SPBs^38, 48^. We next identified proteins that show reduced Sad1-3V5 association in cells lacking Csi1, Lem2 or both. Enrichment of γ-TuRC and kinetochore proteins (Nuf2, Ndc80) with Sad1-3V5 was reduced in *csi1Δ* cells (Figure 1D), whereas levels of ESCRT proteins, dynein light chain, monopolin proteins, and the Lem2-binding protein Nur1/Mug154 were reduced in *lem2Δ* cells (Figure 1E). Both sets of proteins also showed specific reduction in their association with Sad1-3V5 in *csi1*Δ*lem2*Δ cells, with essentially no loss of additional proteins (Figure S1A). These findings are consistent with the known functions of Csi1 in promoting spindle formation and centromere clustering^38, 54^ and the role of Lem2 in nuclear membrane remodeling^55–57^. Although Csi1 clearly mediates association of some kinetochore proteins with Sad1-3V5, enrichment of Mis18C components was unaffected by Csi1 and/or Lem2 loss, suggesting that Mis18C may associate with Sad1-3V5 by a distinct route (Figures 1D-E and S1A-B).

### TEV site insertions allow cleavage and loss of the Sad1 nucleoplasmic region *in vivo*

The domain organization of Sad1 indicates that only the first N-terminal 167 residues (before the transmembrane domain) protrude into the nucleoplasm (Figure 2A). Additionally, the temperature sensitive *sad1-2* mutation (T3S, S52P) causes centromeres to detach from interphase SPBs^51^. SPB-centromere clustering therefore relies on this N-terminal region. To selectively and inducibly disrupt this region, we inserted a tobacco etch virus (TEV) protease cleavage site at amino acid positions 60 or 140 in the 3V5-tagged endogenous *sad1^+^* gene to cleave Sad1-3V5 protein at these positions upon TEV protease expression (Figure 2A). Nuclear-targeted TEV protease expression cleaved approximately 80% of Sad1-TEV140-3V5 in cells (Figures S2A-S2B). Immunolocalization showed that the C-terminal portion of cleaved Sad1-TEV140-3V5 remained colocalized with a SPB marker (Figure S2C). TEV expression rendered *sad1-TEV140-3V5* cells mildly thiabendazole-(TBZ) and temperature-sensitive compared to control cells, reflecting defective centromere, kinetochore, or spindle function. As expected, this sensitivity depended on TEV expression and the presence of TEV sites (Figure S2D-S2E). The N-terminal product of Sad1-TEV140-3V5 cleavage was presumably degraded as it was undetectable when the 3V5 tag was fused to the N-terminus of Sad1 (3V5-Sad1-TEV140, Figure S2F). We therefore only analyzed the stable C-terminal part of Sad1-3V5 in subsequent cleavage experiments. This system allows inducible *in vivo* cleavage of the Sad1 N-terminus and disruption of Sad1 function.

**Figure 2.**
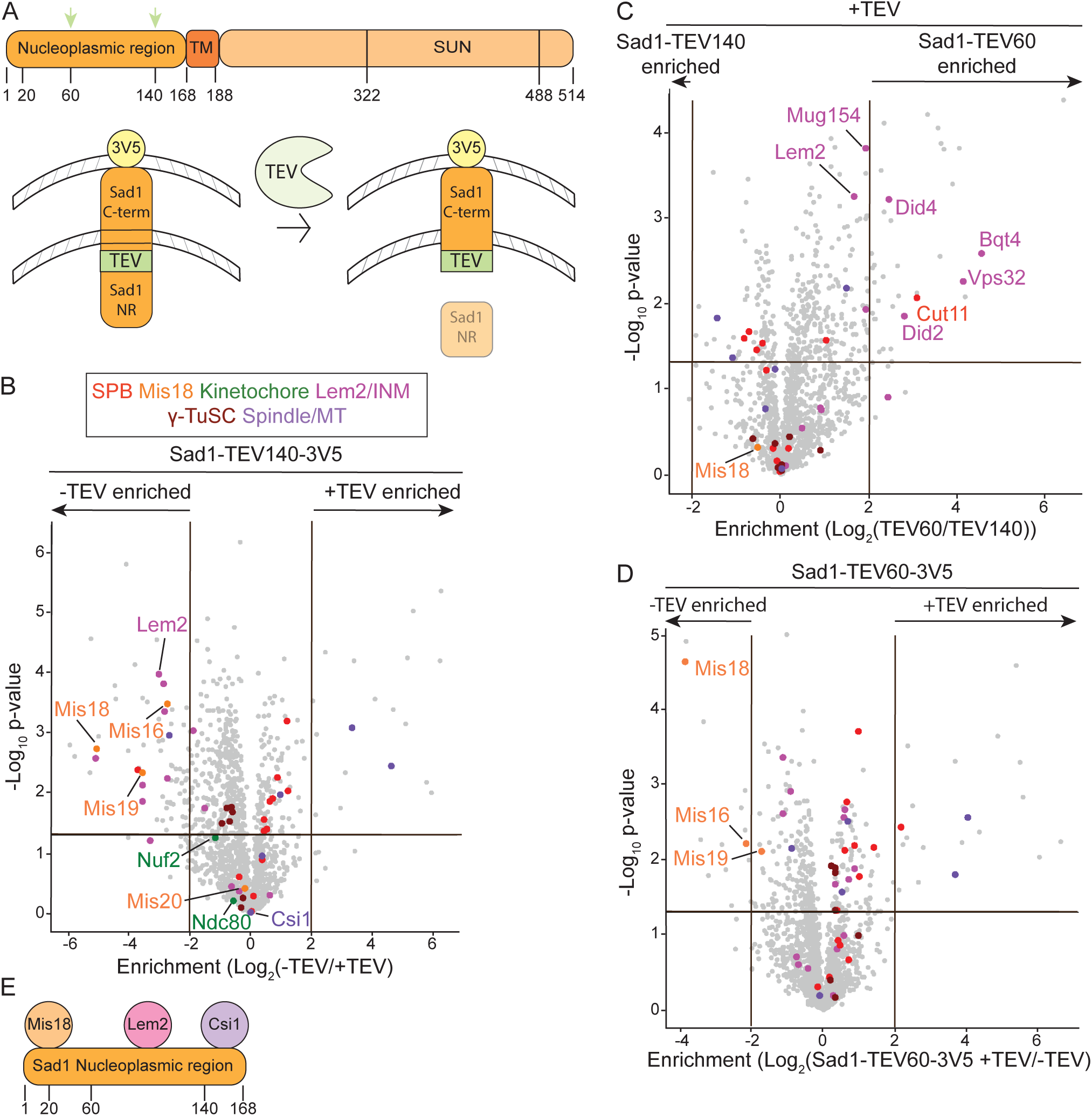
*In vivo* TEV cleavage of the Sad1 N-terminus disrupts Mis18C association with Sad1. (A) Diagram showing Sad1 protein domains and Sad1-TEV cleavage strategy. Top: Sad1 TEV sites were inserted at residues 60 or 140 (green arrows) in the nucleoplasmic region of Sad1. Numbering indicates Sad1 amino acid positions. TM: transmembrane domain. SUN: Sad1/UNC84 lumenal domain. Bottom: Thiamine washout induces nuclear-targeted TEV protease expression (NLS-9myc-TEV-2xNLS), cleaving Sad1 at an inserted TEV recognition site (green). NR: Nucleoplasmic region. (B) Volcano plot of LFQ-MS data comparing proteins enriched in anti-V5 Sad1-TEV140-3V5 IPs before (-) or after (+) TEV cleavage. (C) Volcano plot of LFQ-MS data comparing proteins enriched in anti-V5 Sad1-TEV60-3V5 and Sad1-TEV140-3V5 IPs after TEV cleavage. Dot colors as in (B). (D) Volcano plot of LFQ-MS data comparing proteins enriched in anti-V5 Sad1-TEV60-3V5 IPs before (-) or after (+) TEV cleavage. Dot colors as in (B). (E) Cartoon summarizing Mis18C-, Lem2-and Csi1-interacting regions within the nucleoplasmic region of Sad1 based on LFQ-MS data +/- TEV cleavage.

### Mis18C association with SPBs depends on the N-terminal region of Sad1

To identify proteins that specifically associate with the Sad1 N-terminal region, we compared sets of proteins associated with uncleaved and cleaved Sad1-TEV140-3V5 by IP/LFQ-MS analysis. Comparison of uncleaved Sad1-TEV140-3V5 and Sad1-3V5 showed that copurifying proteins were largely unaffected by TEV site insertion (Figure S2G). Proteins depleted from Sad1-TEV140-3V5 purifications upon its cleavage include Lem2, Lem2-dependent proteins (see Fig 1E), and Mis18C components (Figure 2B; Table S3). In contrast, levels of Csi1 and Csi1-dependent proteins were unaffected (Figure 2B and see Figure 1D), suggesting that they associate with Sad1 through residues 140-167, which remain between the cleavage site and the transmembrane region.

To locate more precisely which parts of the Sad1 N-terminal region recruit specific proteins, we compared protein enrichments in Sad1-TEV60-3V5 and Sad1-TEV140-3V5 purifications. Lem2 and associated proteins were retained following Sad1-TEV60-3V5 cleavage, suggesting that Lem2 associates with Sad1 mainly through residues 60-140 (Figure 2C). In contrast, Mis18C components were lost following TEV cleavage at position 60, indicating that they are recruited by the N-terminal 60 residues of Sad1 and independently of Lem2 (Figure 2D).

We next examined the *in vivo* impact of Sad1 cleavage on the localization SPB/centromere proteins in cells expressing Sad1-TEV140-3V5. In agreement with LFQ-MS analysis, cleavage delocalized Mis18-GFP from SPB-centromere clusters and diminished Lem2-GFP SPB-associated signals, whereas Csi1-GFP was clearly still visible as a focus despite slightly lower SPB-normalized intensities (Figures 3A and S3A-S3B). Thus, Mis18C and the SPB proteins Lem2 and Csi1 associate with Sad1 through distinct subregions of the Sad1 N-terminal region *in vivo*.

**Figure 3.**
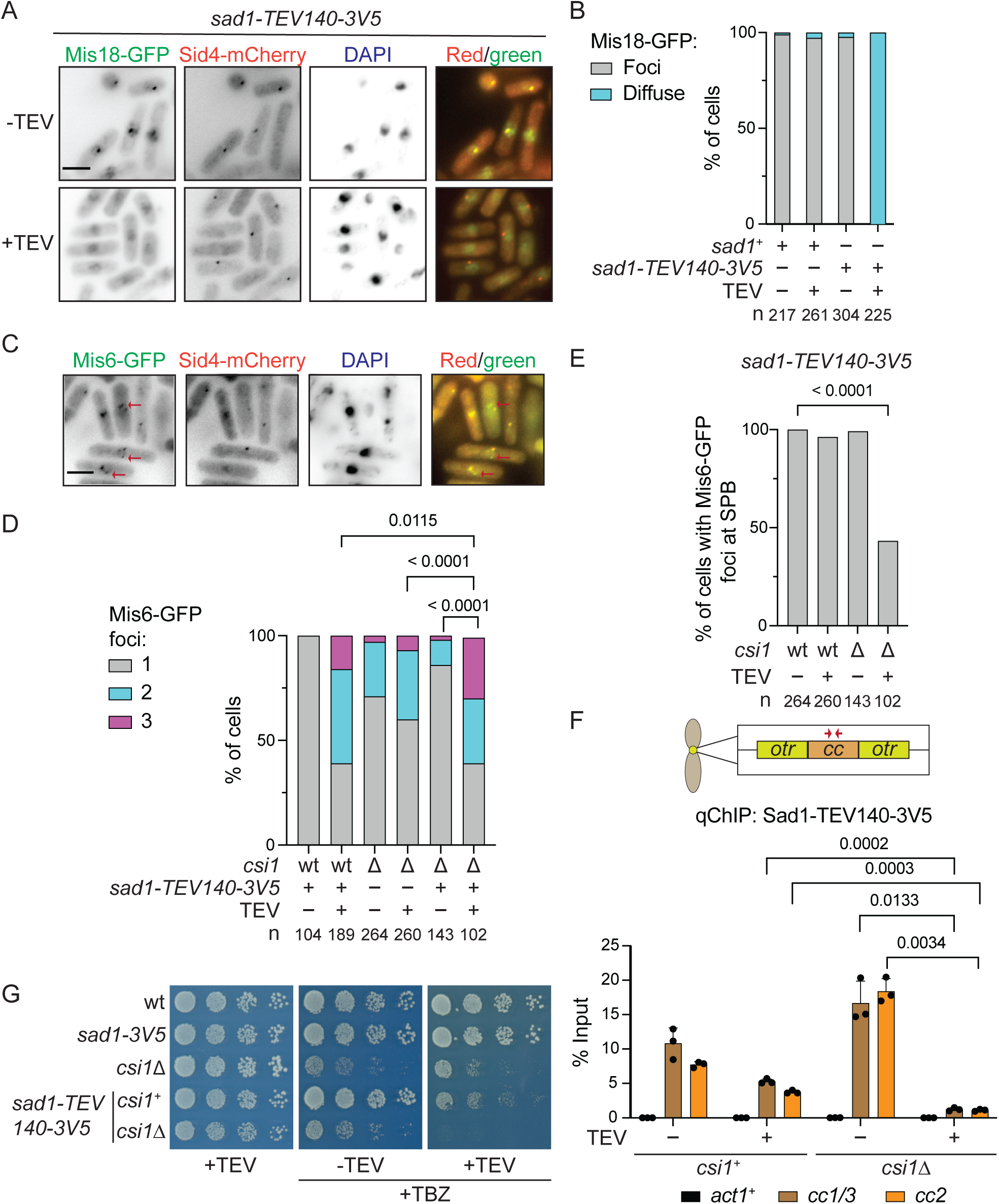
TEV cleavage in the Sad1 nucleoplasmic region dissociates Mis18 and kinetochores from SPBs *in vivo*. (A) Localization of Mis18-GFP and the SPB protein Sid4-mCherry in wild-type *sad1^+^* and *sad1-TEV140-3V5* cells with (+) or without (-) TEV expression induced for 16 hours from *pnmt1-TEV*. (B) Quantification of cells (%, cell number n below) with normal Mis18-GFP focus or diffuse signal. (C) Example localization of Mis6-GFP and SPB protein Sid4-mCherry in *sad1-TEV140-3V5* cells with TEV expression. (D) Quantification of cells (%, cell number n below) with 1, 2 or 3 separate Mis6-GFP foci (arrows). Wild-type *sad1^+^* (wt) or *sad1-TEV140-3V5* cells were grown with (+) or without (-) TEV expression. P-values calculated by χ^2^ test. (E) Quantification of cells (%, cell number n below) retaining a Mis6-GFP focus at SPBs in wild-type (wt) and *csi1Δ* cells with (+) or without (-) Sad1-TEV140-3V5 cleavage. P-value calculated by χ^2^ test. (F) anti-V5 ChIP-qPCR for Sad1-TEV140-3V5 on central core domains of centromeres (*cc1/3* and *cc2*, red arrows) and the control *act1* locus (actin) in wild-type and *csi1*Δ cells with (+) or without (-) Sad1-TEV140-3V5 cleavage. Error bars are SD. P-values calculated with Welch’s t-test. (G) Serial dilution growth assays of wild-type *csi1^+^* and *csi1Δ* cells on plates with (+) or without (-) TEV expression in the presence or absence of TBZ at 10 lg/mL. Wild-type (wt), *sad1-3V5* and *csi1*Δ cells included as controls for TBZ sensitivity.

### SPB-centromere clustering involves multiple interactions via the Sad1 N-terminal region

Sad1 cleavage enabled us to dissect how centromeres associate with SPBs. The Mis6-GFP kinetochore protein marks centromere location and thereby indicates the integrity of SPB-centromere clustering. All three centromeres cluster, producing a single Mis6-GFP signal at G2 SPBs, as quantified in wild-type cells expressing uncleaved Sad1-TEV140-3V5 (Figure 3C and 3D). Cleavage of Sad1-TEV140-3V5 dramatically disrupted SPB-centromere clustering with many cells displaying two or three distinct Mis6-GFP foci (Figure 3C and 3D), similar to cells lacking Csi1^38^. However, as the association of Csi1 with Sad1 remains intact upon Sad1-TEV140-3V5 cleavage (Figure 2B), the declustering phenotype suggested that the Sad1 N-terminal region contributes to two mechanisms of SPB-clustering: one operating through residues 1-140 and the other via Csi1 association with residues 140-167 that remain after TEV cleavage. To test this possibility, we quantified the number of separate Mis6-GFP centromere foci in wild-type and *csi1Δ* cells with and without Sad1-TEV140-3V5 cleavage. The proportion of cells with three Mis6-GFP foci (declustered centromeres) upon Sad1-TEV140-3V5 cleavage was greater in *csi1Δ* than *csi1^+^* cells, and increased relative to *csi1*Δ cells with intact Sad1 (Figure 3D). Strikingly, almost half of *csi1Δ* cells with cleaved Sad1-TEV140-3V5 showed complete dissociation of all Mis6-GFP marked centromeres from SPBs, an arrangement not observed in cells with only Sad1-TEV140-3V5 cleaved or just lacking Csi1 (Figure 3E).

As a distinct approach to determine the impact of Sad1-TEV140-3V5 cleavage, we assessed SPB-centromere association using quantitative chromatin immunoprecipitation (qChIP). Sad1 is enriched over the central domain of *S. pombe* centromeres^38^. We found that Sad1-TEV140-3V5 cleavage significantly reduced Sad1 enrichment on centromere DNA when combined with *csi1*Δ (Figure 3F). An increase in TBZ sensitivity of *csi1*Δ detected when combined with Sad1-TEV140-3V5 cleavage is also consistent with SPB-centromere clustering contributing to kinetochore function during mitosis (Figures 3G and S3C). We conclude that SPB-centromere association via the Sad1 nucleoplasmic region is mediated by two pathways, only one of which requires Csi1 function.

### Purified Sad1 nucleoplasmic region directly binds Mis18

Mis18 appeared to associate with Sad1 independently of centromeres, suggesting that Mis18 may interact directly with Sad1. *In vitro* binding assays with recombinant proteins showed that MBP-Sad1-N (residues 2-167) can bind to GFP-Mis18, but not GFP alone (Figure 4A). In addition, His-Sad1-N and His-Mis18 co-expressed in *E. coli* co-eluted in size-exclusion chromatography, indicating that they form a complex (Figure 4B). Thus, the Sad1 nucleoplasmic region exhibits robust direct binding with Mis18.

**Figure 4.**
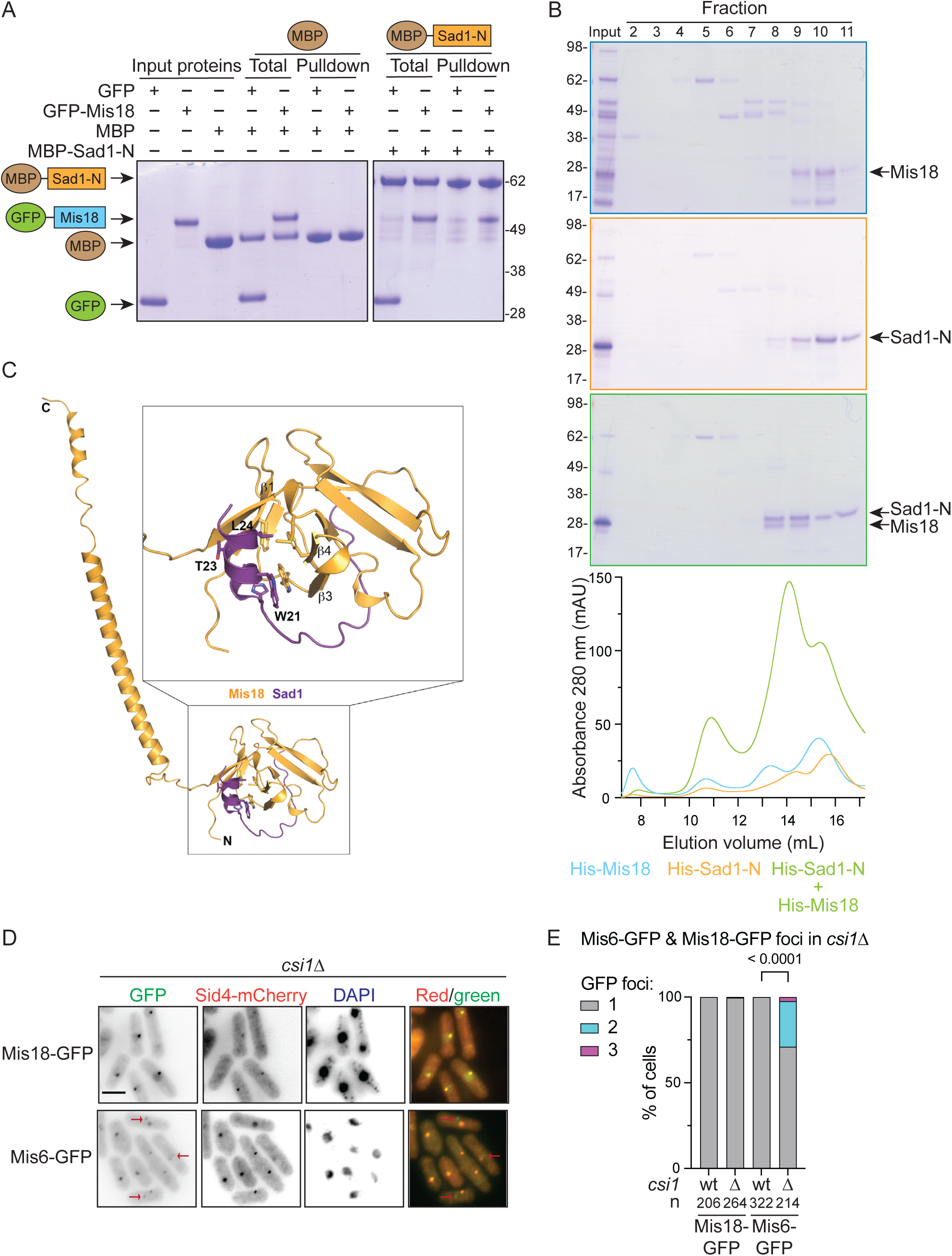
Mis18 directly binds the N-terminal region of Sad1 *in vitro*. (A) *In vitro* binding assays for recombinant GFP-Mis18 full length or GFP (control) to MBP-Sad1-N (residues 2-167, nucleoplasmic region) or MBP bound to amylose resin. Size markers: kDa. (B) His-tagged full length Mis18 and Sad1-N (nucleoplasmic region, 1-167) proteins were expressed individually or coexpressed, affinity purified, and fractionated on a Superdex 200 10/300 GL column. Elution profiles for each protein alone or coexpressed shown below. Size markers: kDa. (C) Mis18 residues 1-194 and Sad1 residues 1-167 were modeled using Colabfold Alphafold2 notebook^74^. Visualization and analysis of interactions were performed using PyMOL. (D) Localization of Mis6-GFP (top) or Mis18-GFP (bottom) and SPB protein Sid4-mCherry in *csi1*⊗ DAPI stained cells (wild-type not shown). Red arrows indicate cells with centromeres (Mis6-GFP) not at SPBs in *csi1*⊗ cells. Scale: 5 μm. (E) Quantification of cells (%, cell number n below) with 1, 2 or 3 Mis6-GFP or Mis18-GFP foci in wild-type (wt) and *csi1*⊗ cells. P-value calculated by χ^2^ test.

Similar *in vitro* binding assays were used to further dissect which regions of Mis18 mediate Sad1-N binding. Neither the N-terminal Yippee domain of Mis18 (residues 1-120), nor the alpha-helical C-terminal region (121-194) exhibited MBP-Sad1-N binding (Figure S4A). Furthermore, removal of the basically charged C-terminal tail from Mis18 (residues 169-194, Mis18_1-168_/Mis18ΔC)^58^ prevented its association with MBP-Sad1-N (Figure S4B). *S. pombe* cells expressing only Mis18ΔC are viable, but temperature sensitive, indicating Mis18 function is compromised (Figure S4C). Moreover, coIP assays showed that the in vivo association of Sad1-3Flag with Mis18ΔC-3V5 *i* was reduced relative to that of full length Mis18-3V5 (Figure S4D).

To gain structural insights into the mode of Sad1 binding to Mis18, we generated an Alphafold model. Our high-confidence model predicts direct interaction between a short Sad1 helical segment spanning amino acid residues 18-25 with a hydrophobic pocket formed by the Mis18 Yippee domain (Figure 4C). Interestingly, the equivalent surface of human Mis18α has been implicated in M18BP1 binding and was shown to be critical for centromere recruitment of human Mis18C^30^.

The *in vitro* binding assays and the difference between Mis18-GFP and Mis6-GFP localization in cells with cleaved Sad1-TEV140-3V5 (diffuse vs foci, respectively, Figure 3A-C) suggest that in *S. pombe* Mis18 is recruited to SPB-centromere clusters primarily via SPBs, rather than via centromeres or constitutively associated kinetochore proteins. This conclusion is supported by our finding that Mis18-GFP localization was unaffected in *csi1Δ* cells whereas Mis6-GFP centromere signals were frequently detected away from SPBs (Figure 4D-4E). Thus, Mis18 G2 localization appears to be mainly driven by its direct association with Sad1 at SPBs, not centromeres.

### Sad1 recruits Mis18 to SPBs independently of centromeres

To determine if centromere clustering with SPBs depends on the association of Sad1 with Mis18, we sought conditions that more precisely disrupt Mis18 recruitment to SPBs. The N-terminal 60 Sad1 residues mediate Mis18C binding (Figures 2B-2E), and our structural analysis predicted that Sad1 interacts with Mis18 through the short alpha helical 21-WSTL-24 region (Figures 4C and S5A). We therefore replaced WSTL with four alanine residues, generating the *sad1-4A* mutation at the endogenous *sad1* locus. *sad1-4A* did not significantly affect Sad1 protein levels, localization, or growth under standard conditions (Figures 5A and S5B-S5D). However, the localization of Mis18-GFP and the Mis18C component Eic1/Mis19-GFP at SPBs was disrupted in *sad1-4A* cells, giving diffuse signals similar to that of Mis18-GFP upon Sad1-TEV140-3V5 cleavage (Figures 5B and S5E). In contrast, Mis6-GFP-marked centromeres and the CENP-A^Cnp1^ chaperone HJURP^Scm3^ remained localized close to SPBs in *sad1-4A* cells (Figures 5B and S5F). Furthermore, *sad1-4A* disrupted Mis18-3Flag coIP with Sad1-3V5 (Figure S5G). Thus, mutation of four residues close to the Sad1 N-terminus disrupts association of Mis18, but not centromeres, with SPBs.

**Figure 5.**
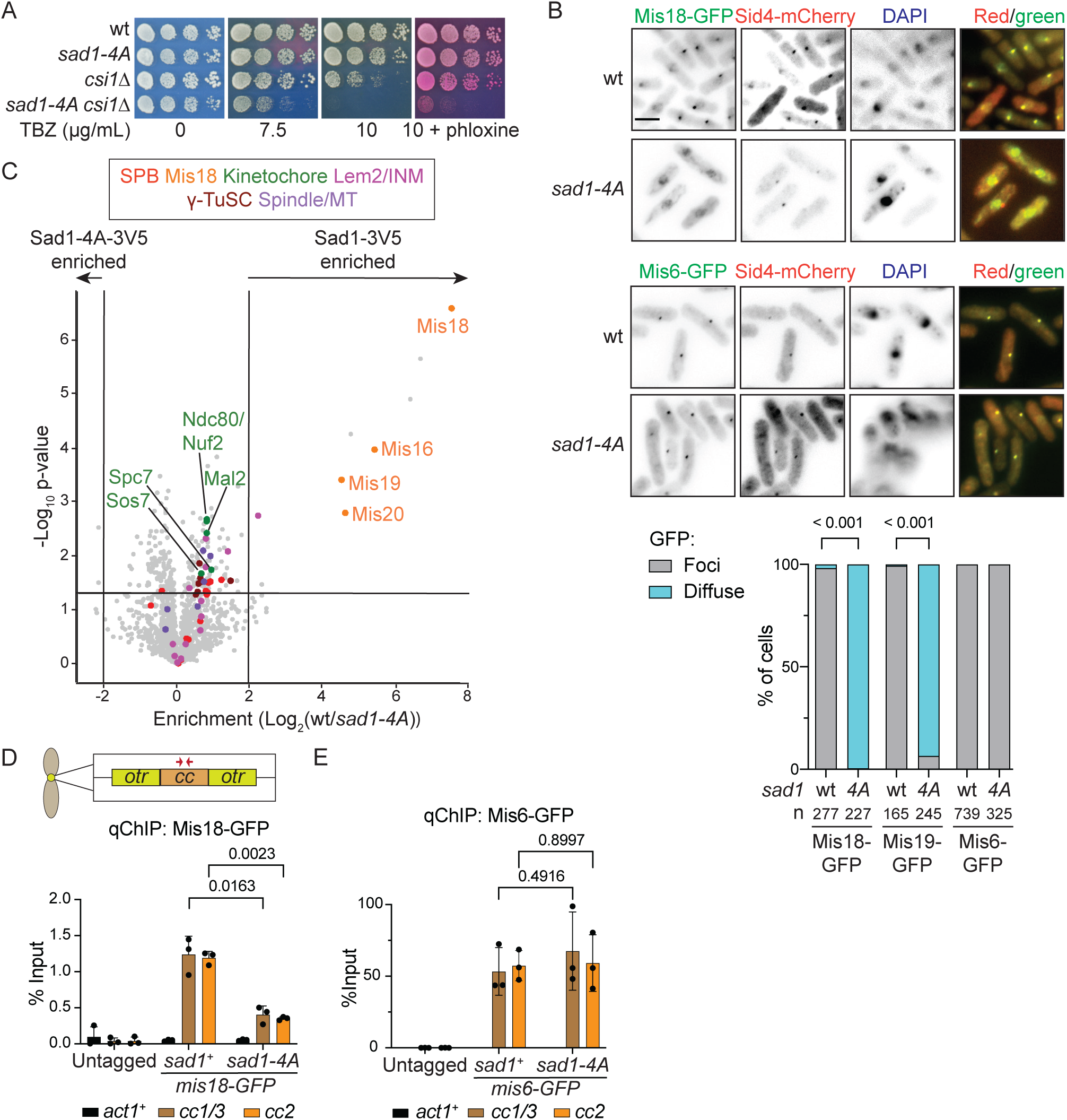
The *sad1-4A* mutation disperses Mis18, but not centromeres, from SPBs. (A) Serial dilution growth assays of wild-type, *sad1-4A*, *csi1*⊗ and *sad1-4A csi1*⊗ cells plated on YES media with or without TBZ added at the indicated concentrations. Phloxine red staining indicates inviable cells. (B) Localization of Mis18-GFP (top) or Mis6-GFP (middle) Mis19-GFP (see Figure S5E) and SPB protein Sid4-mCherry in wild-type (wt) or *sad1-4A* DAPI stained cells. Scale bar: 5 μm. Bottom: Quantification of cells (%, cell number n below) with Mis18-GFP, Mis19-GFP or Mis6-GFP foci or diffuse signals in wild-type (wt) and *sad1-4A* cells. P-values calculated by χ^2^ test. (C) Volcano plot of LFQ-MS data comparing proteins enriched in anti-V5 Sad1-4A-3V5 and Sad1-3V5 triplicate IPs. (D, E) anti-GFP ChIP-qPCR for Mis18-GFP (D) and Mis6-GFP (E) on central core domains of centromeres (*cc1/3* and *cc2*, red arrows) and *act1* control locus (actin) in *sad1^+^* wild-type and *sad1-4A* cells and wild-type cells with no GFP (Untagged). Bar plots show mean values with standard deviation. P-values calculated with Welch’s t-test.

To assess the specificity of *sad1-4A* for disrupting the Mis18-Sad1 interaction, we compared proteins enriched in Sad1-3V5 and Sad1-4A-3V5 IPs. LFQ-MS analyses showed that almost exclusively Mis18C components were depleted in Sad1-4A-3V5 purifications (Figure 5C, Table S4). In addition, qChIP revealed that enrichment of Mis18-GFP at centromeres was significantly reduced in *sad1-4A* cells, whereas Mis6-GFP enrichment was unaffected (Figures 5D-5E). The Sad1-4A mutant protein is unable to directly recruit Mis18C to SPBs but does not detectably perturb centromere clustering at SPBs. Nonetheless, genetic analysis showed that, as with Sad1-TEV140-3V5 cleavage, the *sad1-4A* mutation enhanced the TBZ sensitivity of cells when combined with *csi1* or *bub1* gene deletions, both of which have roles in chromosome segregation (Figures 5A and S5H-S5I). However, deletion of *lem2* showed no genetic interaction with *sad1-4A* (Figure S5I). Such phenotypes are consistent with compromised centromere function in *sad1-4A* cells and suggest that direct Sad1-Mis18 binding contributes to functions at centromeres that are distinct from SPB-centromere clustering.

### Sad1-Mis18 association promotes CENP-A^Cnp1^ chromatin establishment and maintenance

A key function of Mis18C is to direct CENP-A^Cnp1^ deposition and thus its maintenance at centromeres^2^. We therefore tested if CENP-A^Cnp1^ incorporation at endogenous centromeres is compromised when Mis18 association with Sad1 is disrupted. However, qChIP revealed no reduction in CENP-A^Cnp1^ levels on the central domain of *cen2* (*cc2*) in *sad1-4A* cells relative to wild-type (Figure 6A).

**Figure 6.**
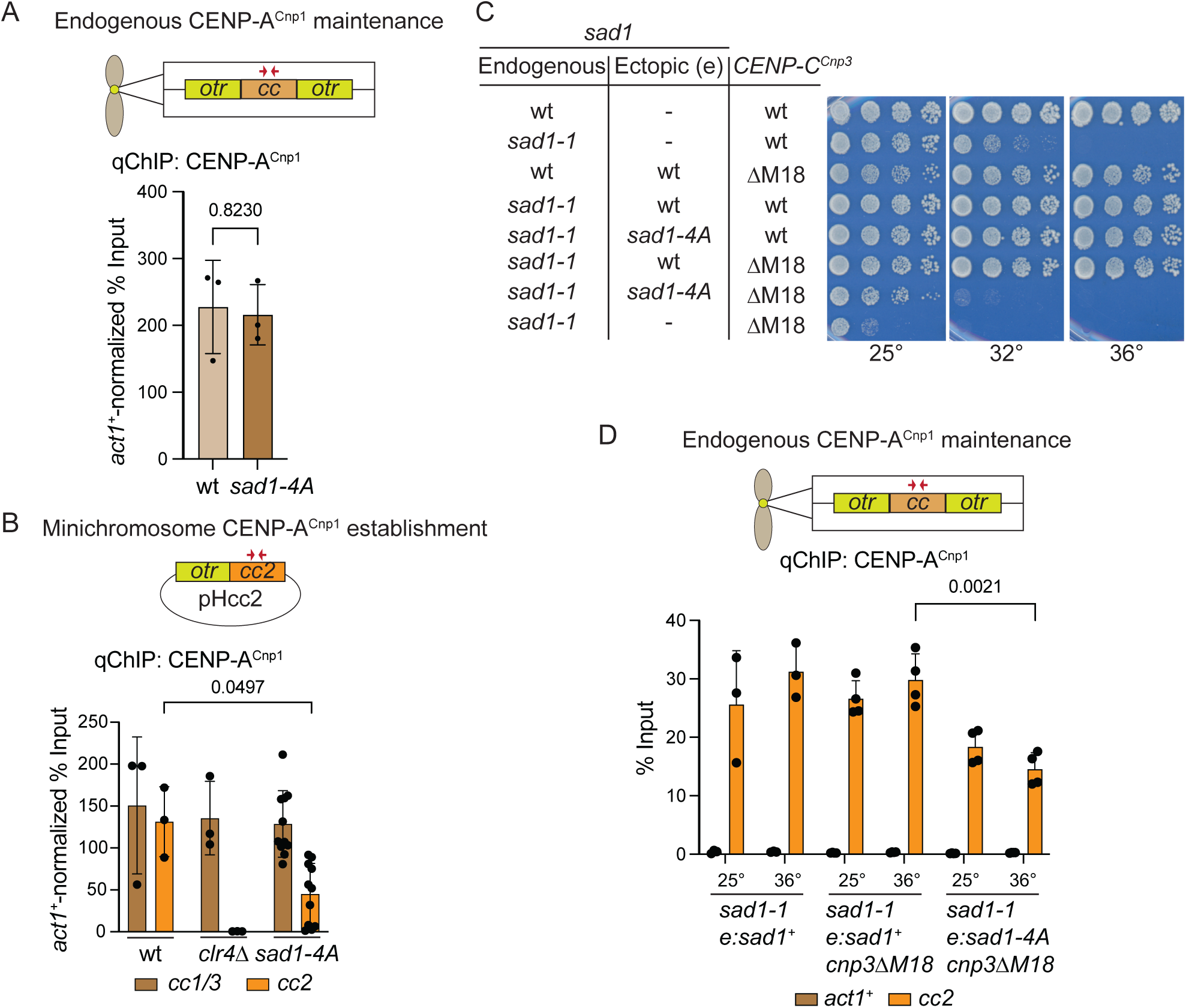
Mis18 utilizes Sad1 and CENP-C^Cnp3^ for CENP-A^Cnp1^ incorporation at centromeres. (A) Quantitative anti-CENP-A^Cnp1^ ChIP-qPCR for CENP-A^Cnp1^ on central core domains of centromeres (*cc1/3* and *cc2*, red arrows) in wild-type and *sad1-4A* cells normalized relative to *act1* locus (actin). (B) pHcc2 minichromosome DNA was transformed into wild-type, *clr4*⊗ and *sad1-4A* cells. anti-CENP-A^Cnp1^ ChIP-qPCR was performed to assess the levels of CENP-A^Cnp1^ chromatin established on the central *cc2* domain (red arrows) and control endogenous centromeric *cc1/3* loci in multiple individual pHcc2 transformants relative to CENP-A^Cnp1^ on the *act1* control locus. *cc2* is unique to pHcc2 in these cells because endogenous *cc2* is replaced with *cc1*^59^. (C) Serial dilution growth assays of the indicated strains plated on YES media at 25°C, 32°C, or 36°C. (D) anti-CENP-A^Cnp1^ ChIP-qPCR performed to assess the level of CENP-A^Cnp1^ chromatin on the endogenous central *cc2* domain of *cen2* (red arrows) relative to the *act1* control locus in cells with the temperature sensitive *sad1-1* mutation covered by ectopically integrated *sad1^+^* or *sad1-4A* genes. Error bars: SD. P-values: Welch’s t-test.

In addition to CENP-A^Cnp1^ maintenance at endogenous centromeres, Mis18C may also be required to establish CENP-A^Cnp1^ chromatin on naïve centromere DNA templates. We therefore used a minichromosome-based establishment assay to test if SPB-associated Mis18 is required for *de novo* CENP-A^Cnp1^ chromatin assembly^59^. pHcc2 minichromosome DNA, bearing a central *cc2* core and flanking outer repeat, efficiently incorporated CENP-A^Cnp1^ on *cc2* upon transformation into wild-type but not *clr4*Δ negative control cells (Figure 6B). In *sad1-4A* cells, the levels of CENP-A^Cnp1^ chromatin assembled on *cc2* in pHcc2 transformants was significantly reduced relative to wild-type. There was considerable variability in CENP-A^Cnp1^ levels incorporated across 11 independent transformants, indicating that *de novo* CENP-A^Cnp1^ establishment was reduced but not eliminated in *sad1-4A* cells. Thus, although the SPB-Sad1-associated Mis18 pool is not required to maintain existing CENP-A^Cnp1^ at endogenous centromeres, it ensures that CENP-A^Cnp1^ chromatin can be efficiently established when CENP-A^Cnp1^ and kinetochore proteins are absent from substrate DNA.

*Sad1-4A* cells are viable, despite the loss of Mis18 from SPBs, whereas Mis18 is essential for viability^21^. Moreover, the retention of some Mis18 at centromeres in *sad1-4A* cells (Figure 5E) suggests the existence of a separate centromere-based Mis18 recruitment pathway that may permit CENP-A^Cnp1^ maintenance and *sad1-4A* viability when the main SPB-Sad1-associated Mis18 pool is dispersed. *S. pombe* Mis18 is known to bind CENP-C^Cnp3^ and disruption of this interaction in *cnp3ΔM18* (residues 325-490 removed) mutant cells results in slow growth and TBZ sensitivity (Figure S6A-S6B) and reduced CENP-A^Cnp1^ signals at centromeres^31^. Growth of *cnp3ΔM18* cells was also inhibited when Mis18C was released from SPBs by Sad1-TEV60-3V5 cleavage (Figure S6A). Moreover, genetic analysis demonstrated that *sad4-A cnp3ΔM18* double mutant cells are not viable (Figure S6C). These genetic interactions are consistent with the operation of distinct SPB-Sad1- and centromere-CENP-C^Cnp3^-based pathways for Mis18C recruitment to SPB-centromere clusters.

We therefore used the *cnp3ΔM18* mutation to test if disruption of CENP-C^Cnp3^-Mis18 association renders CENP-A^Cnp1^ chromatin assembly dependent upon the main Sad1-associated Mis18 pool. *cnp3ΔM18* cells bearing the *sad1-1* temperature sensitive mutation were constructed with the *sad1* gene promoter driving either wild-type Sad1 or Sad1-4A protein expression from an ectopic locus (*e*). Shifting these cells to 36°C inactivates Sad1-1, allowing the phenotype of *e:sad4-A cnp3ΔM18* double mutants to be assessed relative to e:*sad1^+^ cnp3ΔM18* cells. As expected, *sad1-1 e:sad1^+^ cnp3ΔM18* cells were viable at both permissive (25°C) and restrictive (36°C) temperatures. In contrast, *sad1-1 e:sad4-A cnp3ΔM18* were inviable at 36°C even though Sad1-4A was expressed at similar levels to wild-type Sad1 (Figures 6C and S6D). qChIP revealed that the levels of CENP-A^Cnp1^ on endogenous centromeres was significantly reduced in *sad1-1* e:*sad1-4A cnp3ΔM18* relative to *sad1-1* e:*sad1^+^ cnp3ΔM18* cells at 36°C (Figure 6D). Thus, the *sad4-A* and *cnp3ΔM18* mutations act synergistically to impair CENP-A^Cnp1^ incorporation. We conclude that two pathways operate to efficiently recruit Mis18 to centromeres and ensure maintenance of normal CENP-A^Cnp1^ levels (Figure 7): one through direct association of Mis18 with Sad1 at SPBs and the other via CENP-C^Cnp3^ at centromeres.

**Figure 7.**
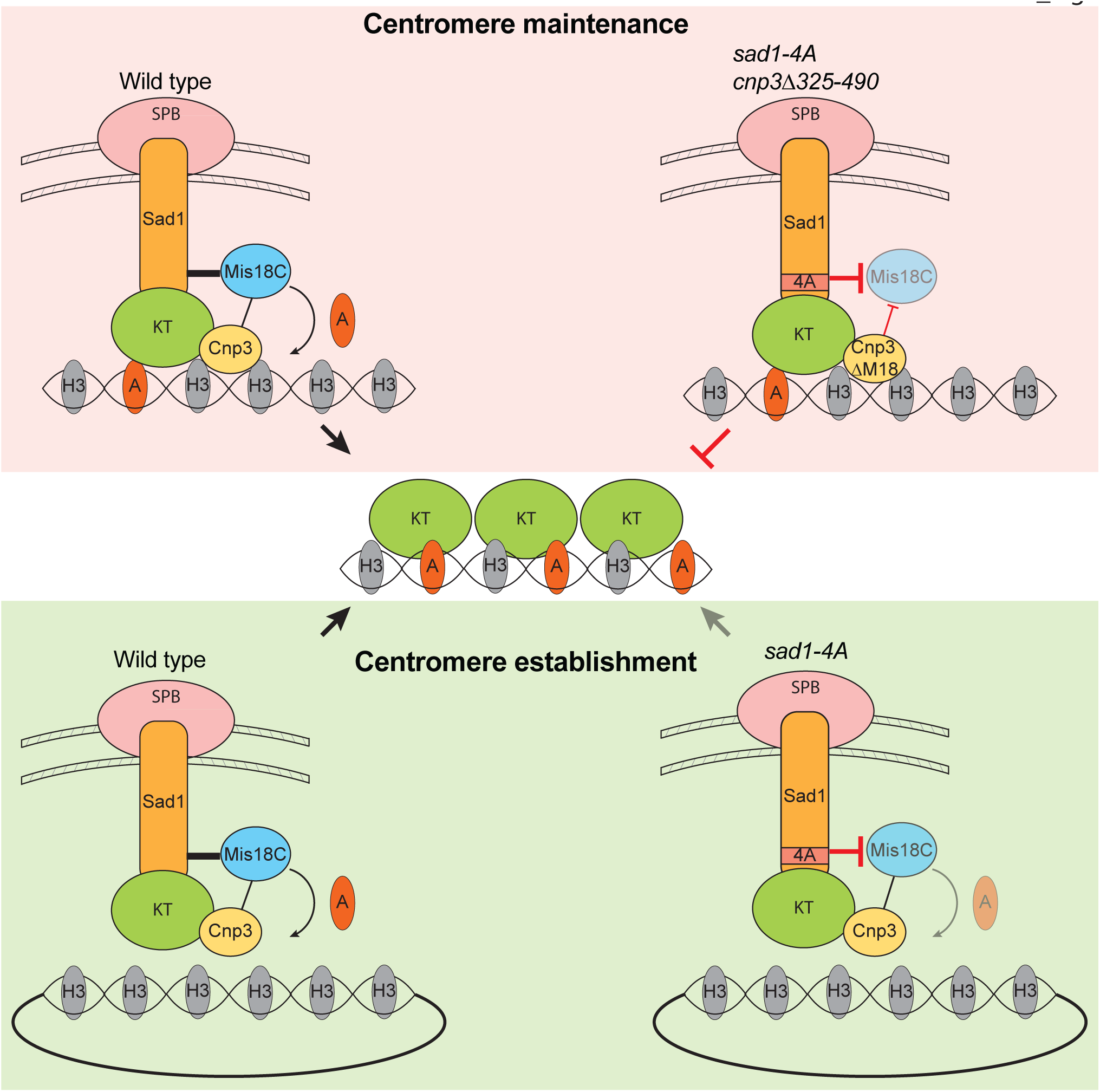
Model showing parallel Sad1 and CENP-C^Cnp3^ pathways of Mis18 recruitment and CENP-A^Cnp1^ incorporation. Endogenous centromeres are maintained by Mis18C recruited to SPBs through Sad1 and to centromeres through CENP-C^Cnp3^. The bulk of Mis18C is localized through the Sad1 interaction (thick line) and a small proportion of Mis18C is recruited to centromeres through CENP-C^Cnp3^ (thin line). Abolishing Mis18C recruitment with *sad1-4A* and *cnp3ΔM18* inhibits CENP-A^Cnp1^ assembly at engeonous centromeres, causing lethality. *De novo* centromere formation on minichromosomes that contain appropriate sequence undergo CENP-A^Cnp1^ establishment when they are in close proximity to SPBs and therefore Mis18C. In *sad1-4A*, Mis18C at the SPB is diminished leading to reduced *de novo* CENP-A deposition on minichromosomes (light arrow).

## Discussion

Mis18C is required to maintain CENP-A chromatin, and thus kinetochore integrity, by targeting the CENP-A chaperone HJURP to pre-existing CENP-A chromatin. To understand how centromeres are established and propagated, and how neocentromere formation is suppressed, it is important to determine how Mis18C is recruited to its chromatin substrate. In mammalian cells, Mis18α/β are recruited to pre-existing CENP-A chromatin by binding the adaptor protein M18BP1 which associates with CENP-C at existing centromeres. However, various eukaryotes, including *S. pombe* and *D. melanogaster*, lack a M18BP1 ortholog, suggesting that other mechanisms of Mis18 recruitment are of primary importance in these systems and that similar mechanisms may also contribute to CENP-A chromatin maintenance even in species exhibiting the canonical Mis18-M18BP1 pathway^25, 60–63^. In many organisms centromeres tend to cluster, a phenomenon that is particularly prevalent in various fungi and plants where centromeres cluster near SPBs^33–35, 64, 65^. Here, we demonstrate that in fission yeast Mis18C is directly recruited to SPBs, thereby concentrating it near centromeres that robustly cluster near SPBs in G2 cells when CENP-A^Cnp1^ is replenished. Our analyses show that this Mis18-SPB pathway is needed for efficient CENP-A chromatin establishment on centromere DNA and also acts in parallel to a conventional Mis18-CENP-C^Cnp3^ pathway to preserve CENP-A^Cnp1^ chromatin at centromeres.

Our TEV-induced cleavage strategy identified proteins that specifically associate with the N-terminal nucleoplasmic region of Sad1, providing insight into the constituents on the nucleoplasmic side of *S. pombe* SPBs (Figures 1 and 2). Components of Mis18C were found to associate with Sad1 via its N-terminal 60 residues. Mis18C association with Sad1 involves direct binding of Mis18 with Sad1 and this interaction was disrupted by the specific *sad1-4A* N-terminal mutation. In contrast, the Lem2 and Csi1 proteins, that are known to contribute to SPB-centromere clustering, appeared to associate with Sad1 through residues 60-140 and 140-167, respectively. This modular arrangement suggests that Mis18C recruitment and SPB-centromere clustering are separable Sad1 functions. Indeed, the *sad1-4A* mutation dispersed most Mis18-GFP away from SPBs without affecting centromere (Mis6-GFP) clustering at SPBs (Figure 5B). Reciprocally, most Mis18-GFP remained at SPBs and did not colocalize with centromeres which had separated from G2 SPBs in *csi1*Δ cells (Figure 4E). Thus, the congregation of centromeres at SPBs and Mis18C recruitment to SPB-centromere clusters are mediated by distinct segments of the Sad1 nucleoplasmic region.

Sad1 is located at SPBs throughout the cell cycle, whereas Mis18C is released from the SPB-centromere cluster upon mitotic entry^21, 24, 47^, suggesting that association of Mis18 with Sad1 might be regulated by post-translational modification. *S. pombe* Cdk1 (Cdc2) and Polo (Plo1) kinases are active at SPBs upon mitotic onset, just after maximal CENP-A^Cnp1^ incorporation in late G2, when Mis18 is released from SPBs^17, 18, 66^. Mitotic phosphorylation of both Sad1 and Mis18 have been reported^67, 68^ and Plo1 kinase activity is required to reconfigure Sad1 into a ring around SPBs at the end of G2^69^. Since unphosphorylated recombinant Mis18 directly associates with the Sad1 N-terminal region *in vitro* (Figures 4A-4B), we surmise that phosphorylation of Sad1 and/or Mis18 may disrupt their association to coordinate release of Mis18C from SPBs with disengagement of centromeres, mitotic entry and spindle formation. Interestingly, the association of Mis18α with the M18BP1 adapter protein in human cells has been shown to be regulated by Polo and Cdk1 kinase-mediated phosphorylation - promoting and preventing their association, respectively^26, 70–73^.

Association of the conserved Yippee domain of hMis18α with hM18BP1 is required for hMis18α recruitment to human centromeres^25, 26, 30, 72^. As our structural analyses predict that the Yippee domain of *S. pombe* Mis18 mediates its interaction with the Sad1 N-terminal region (Figure 4C), we conclude that Sad1 performs the equivalent function of human M18BP1 in mediating *S. pombe* Mis18C recruitment^26, 27, 30, 72^. Our finding that neither the N- nor C-terminal domains of Mis18 were sufficient to bind the Sad1 nucleoplasmic region *in vitro* (Figure S4A), and that the short Mis18 C-terminal tail was required for robust binding (Figures S4B-S4D), suggests additional binding interactions between Mis18 and Sad1 may exist.

Because Mis18 is required for the assembly of CENP-A^Cnp1^ chromatin^21^, our discovery of a direct Mis18-Sad1 interaction suggested that *S. pombe* SPBs might provide a platform to ensure CENP-A^Cnp1^ replenishment during G2. Disruption of either the Mis18-Sad1 interaction (*sad1-4A;* Figure 6A) or Mis18 recruitment to centromeres via CENP-C^Cnp3^ (*cnp3ΔM18*; Figure 6D)^31^ did not impact CENP-A^Cnp1^ levels at endogenous centromeres. However, combining *sad1-4A* and *cnp3ΔM18* in conditionally mutant cells resulted in lower CENP-A^Cnp1^ levels at centromeres and lethality (Figures 6C-6D). These observations indicate that the SPB-Sad1-Mis18 pathway identified here operates redundantly with a centromere-based CENP-C^Cnp3^-Mis18 pathway to maintain CENP-A^Cnp1^ chromatin. We also observed significantly lower levels of CENP-A^Cnp1^ chromatin establishment on naïve minichromosome-borne centromere DNA (Figure 6B) in *sad1-4A* cells, demonstrating that Mis18-Sad1 association is required to promote robust *de novo* CENP-A^Cnp1^ chromatin assembly (Figure 7).

*S. pombe* pericentromeric heterochromatin appears to localize the central core domain of centromeres to the nuclear periphery, where it can encounter existing centromeres at SPBs and promote efficient *de novo* CENP-A^Cnp1^ chromatin assembly^41, 42, 59^. Moreover, inserting a naïve centromeric DNA template near an existing centromere or tethering a plasmid with naïve centromeric DNA to SPBs promotes CENP-A^Cnp1^ establishment independently of flanking heterochomatin^41^. These findings led us to propose that the centromere cluster provides a microenvironment where CENP-A^Cnp1^ assembly factors (Mis18C and HJURP^Scm3^) are concentrated, promoting CENP-A^Cnp1^ chromatin establishment and maintenance. However, unexpectedly we find that the N-terminal region of Sad1 recruits Mis18C, independently of centromeres or their clustering, thereby providing two factor authentication in centromere specification: naïve centromeric DNA exhibiting appropriate properties and positioned in the right nuclear location (near SPBs) establish CENP-A^Cnp1^ chromatin more efficiently.

Interestingly, centromeres cluster at the nuclear periphery in normal human tissues and this clustering decreases in cancer cells^10^, which frequently also exhibit aberrant CENP-A expression and centromere locations. The Mis18-Sad1-SPB recruitment mechanism we have uncovered here reveals how centromere identity is ensured through the spatial positioning of centromeres in nuclei. It is possible that centromere clustering in human and other cell types also contributes to their establishment and maintenance.

## Acknowledgements

We thank members of team Allshire for input during the course of this work, especially Alison L. Pidoux for suggesting the TEV cleavage strategy and insightful comments on the manuscript. We thank Dave Kelly and Toni McHugh for training on, and maintenance of, our microscopes, Ken Sawin for anti-Cdc11 antibody, the Marston lab for plasmid carrying the TEV protease, Mitsuhiro Yanagida for *mis16-GFP* and *mis18-GFP* strains, Sigurd Braun for the *lem2*Δ strain, and Iain Hagan for providing Keith Gull’s Tat1 antibody. The Edinburgh Protein Production Facility provided protein purification equipment (Multi-User Equipment grant 101527/Z/13/Z).

This work was funded by an HHMI Life Sciences Research Foundation Award to NL, consecutive Wellcome Principal Research Fellowships to R.C.A. supporting N.L. (224358, 200885, 224358), a Wellcome Senior Research Fellowship to A.A.J. supporting B.M. (202811), a Wellcome Instrument grant to J.R. (108504), and core funding for the Wellcome Centre for Cell Biology (203149) supporting C.S.

## Author Contributions

N.L. and R.C.A. conceived the study. Experiments were performed by N.L. B.M. provided recombinant proteins. B.M. and J.P. contributed to experimental design. C.S. performed mass spectrometry and initial data processing facilitated by JR. NL and R.C.A wrote the manuscript.

## Materials and Experimental Procedures

### Yeast Strain construction

*S. pombe* strains used in this study are listed in Tables S5 and S6. C-terminal taggings and deletions were performed on genes at the endogenous locus using standard homologous recombination-based methods^75^. Candidate transformants were verirfied by genomic PCR, microscopy, western blotting, or phenotypic analysis as appropriate. CRISPR/Cas9 was used to insert TEV sites, to N-terminally tag *sad1*, and to generate the *sad1-1* mutation (A323V) at the endogenous *sad1* locus. Cells were co-transformed with the guide RNA plasmid and homologous recombination templates generated by primer annealing, followed by selection and screening^76^. Sad1 N-terminal tagging, TEV-site insertion, and 4A mutations were confirmed by genomic DNA amplification and sequencing with NL266-NL284, and *sad1-1* was similarly verified with NL265-267. *Mis18Δ169-194* and c*np3Δ325-490* were generated by CRISPR-Cas9 and PCR-based verification with NL142-NL360 and NL386-NL387, respectively.

*NLS-9myc-TEV-2xNLS* and ectopic *sad1* alleles were integrated at the *ars1* locus by transforming MluI-digested pNL142 (TEV), BlpI-digested pNL190 (*psad1-sad1*), or BlpI-digested pNL197 (*psad1-sad1-4A)*. Transformants were passaged without selection to single colonies for multiple generations to reduce the likelihood of retaining any episomal DNA and were subsequently crossed to generate the experimental strains. *TEV* and ectopic *sad1* integration were verified by PCR with NL362-NL363. *csi1*Δ was obtained from the Bioneer Deletion Set V1-17E7.

### Yeast growth conditions

TEV expression was induced by thiamine washout. Starter cultures were grown in YES (Yeast Extract with Supplements) media, then washed in PMG (*Pombe* Minimal Glutamate) media without thiamine and used to inoculate sample cultures in PMG supplemented with amino acids and with or without 15 μM thiamine. These cells were grown for 10 hours, or the indicated time, at 32°C. 4x concentrated YES media was used for MS experiments not involving TEV expression. For other experiments, cells were cultured at 32°C in YES. For temperature shifts, starter cultures were grown at 25°C then diluted to approrpiate densities and grown at 25°C or 36°C for 8 hrs. In all cases, cells were cultured to approximate densities of OD_600_ = 0.5-1.0 for microscopy and qChIP experiments or 1.0-2.0 for IP/MS experiments. For growth assays, 1:5 serial dilutions were plated and grown for 3-5 days under the indicated conditions until colonies were fully developed. Phloxine B was used at 2.5 μg/mL.

### Cloning

The TEV-protease expression plasmid was generated by PCR amplifying NLS-9myc-TEV-2xNLS from AMp1325 with NL332-NL333 and inserting into BamHI/SalI-digested pRep1 downstream of the thiamine-repressible *nmt1* promoter^77^ via Gibson assembly. CRISPR/Cas9 targeting constructs were generated by Golden Gate assembly (NEB cat #E1601S) using pLSB plasmids as described^76^. Briefly, target sequence primers were annealed and incubated with the destination vector in the presence of Golden Gate assembly mix at 37°C for 1 hr, then 60°C for 5 min before transformation into *E. coli*. The ectopic *sad1* integration construct (pNL190) was generated by PCR amplifying the *sad1* ORF from genomic *S. pombe* DNA with NL456-NL457, annealing NL458-NL459 to generate the 3Flag sequence, and Gibson assembling these fragments with BamHI-HF/SphI-digested vector (derived from pRad11, a gift from Y. Watanabe). PCR-based mutagenesis of pNL190 was performed with NL453-NL467 to generate the ectopic *sad1-4A* construct (pNL197).

Mis18 codon optimized sequences (GeneArt) were cloned into pET His6 msfGFP TEV (9GFP) cloning vector with BioBrick polycistronic restriction sites or into pEC-K-3C-His (a kind gift from Elena Conti). *Sad1*_2-167_ was amplified with NL389-NL390 and ligated into pET-6His-TEV (9B) or pET-6His-MBP-TEV (9C) with ligation-independent cloning (LIC) to generate 6His-TEV-Sad1_2-167_ or 6His-MBP-Sad1_2-167_ plasmids, repsectively. For the 6His-Sad1_2-167_ 6His-Mis18_FL_ coexpression plasmid, Sad1_2-167_ sequence was excised from pNL188 with NcoI/PacI digestion and ligated with NotI/AsiSI-digested 6His-Mis18 vector. 6HIS-GFP was generated by amplifying GFP with NL461-NL462 and ligating into 9B by LIC. 9B, 9C, and 9GFP were gifts from Scott Gradia. (9B: Addgene plasmid # 48284; http://n2t.net/addgene:48284; RRID:Addgene_48284; 9C: Addgene plasmid # 48286; http://n2t.net/addgene:48286; RRID:Addgene_48286; 9GFP: Addgene plasmid # 48287; http://n2t.net/addgene:48287; RRID:Addgene_48287)

### Protein expression

Analysis of sad1-TEV protein cleavage, TEV protease expression, sad1-4A protein levels, and ectopic sad1 protein levels were performed by lysing cells in Laemmli sample buffer (2% SDS, 10% glycerol, 62.5 mM Tris-HCl pH 6.8, 0.002% bromphenol blue) with 2 mM PMSF and 1 mM DTT by bead beating. Samples were heated to 95°C for 3 min, pelleted at 13.2k RPM in a 4°C microfuge for 5 min, and analyzed by western blotting. α-V5 (Bio-Rad cat #MCA1360) was used at 1:5,000, α-GFP (JL-8, Living Colors cat #632380) was used at 1:1,000, α-Flag (Sigma cat #F1804-5MG) was used at 1:1,000, α-myc (9E10) was used at 1:5,000, and Tat1 (α-tubulin, from Iain Hagan) was used at 1:10,000.

### Protein purification and size exclusion chromatography

6His-Sad1_2-167_, 6His-MBP-Sad1_2-167_, 6His-Mis18, 6His-GFP, and 6His-MBP expression plasmids were transformed into BL21-CodonPlus (DE3)-RIPL *E. coli* (Stratagene, #230280) and recovered on non-inducing MDAG media^78^ with carbenicillin, streptomycin, and chloramphenicol selection. Transformed cells were grown in 2xYT (1.6% tryptone, 1.0% yeast extract, 0.5% NaCl at pH 7.0) with carbenicillin and streptomycin to mid-log phase (O.D._600_ of 0.5-1.0) at 36°C, then induced with 0.1 mM IPTG and cultured at 18°C for 16 hrs. Cells were then pelleted and flash frozen. Cells were resuspended in lysis buffer (20 mM HEPES pH 7.0, 500 mM NaCl, 5% glycerol, 2 mM ditihothreitol (DTT), 1 mM phenylmethylsulfonyl fluoride (PMSF)) and lysed by sonication. Lysate was pelleted at 15,000 x g for 30 min at 4°C. Supernatant was incubated with 1 column volume (CV) of TALON resin (Takara #635502) at 4°C for 1-2 hours. Resin was washed in 20 CV wash buffer (lysis buffer + 5 mM imidazole), then eluted in 0.5-1 mL fractions with elution buffer (20 mM HEPES pH 7.0, 300 mM NaCl, 5% glycerol, 150 mM imidazole, 2 mM DTT). For size exclusion chromatography (SEC) of 6His-Sad1_2-167_ and 6His-Mis18, proteins were concentrated using Amicon 10k MWCO filters (Millipore #UFC901024). SEC was performed with a Superdex 200 10/300 GL column on an ÄKTA Pure^TM^ 25 (Cytiva) system and a buffer consisting of 20 mM HEPES pH 7.0, 150 mM NaCl, and 2 mM DTT.

All other Mis18 constructs were expressed in BL21 Gold cells. 6His-msfGFP-TEV-Mis18_121-194_ and 6His-msfGFP-TEV-Mis18_FL_ were grown in 2xYT while 6His-3C-Mis18_1-120_ and 6His-3C-Mis18_1-168_ were grown in Super Broth. For Mis18_FL_, Mis18_121-194_ and Mis18_1-168_, after entering log phase (O.D._600_ = 0.6-0.8) the temperature was reduce to 18°C for one hour and IPTG was added to final concentration of 0.3 mM and protein was induced overnight. For Mis18_1-120_, induction occured at 25°C for 6 hours.

6His-Mis18_1-168_ was lysed in lysis buffer containing 20 mM Tris (pH8.0 at 4°C), 50 mM NaCl, 35 mM imidazole and 2 mM β-mercaptoethanol (βME) and supplemented with DNase (Sigma #DN25-100mg) to a concentration of 10 µg/ml and cOmplete^TM^ EDTA-free protease inhibitors (Sigma #05056489001), 1 tab per 50 ml. Cells were lysed by sonication and centrifuged at 22,000 rpm for 50 mins at 4°C. Clarified lysates were incubated with HisPur^TM^ Ni-NTA resin (Thermo Fisher #88222) and washed extensively with 100 CV of lysis buffer before elution with lysis buffer containing 500 mM imidazole. The protein was concentrated and SEC was preformed on an ÄKTA system (Cytiva) using a S200 Hiload 16/600 column (Cytiva) equilibrated with 20 mM Tris (pH8.0 at 4°C), 50 mM NaCl, and 4 mM DTT.

6His-Mis18_1-120_ was lysed in lysis buffer containing 20 mM Tris (pH 8.0 at 4°C), 500 mM NaCl, 35 mM imidazole and 5 mM βME and supplemented with DNase to a concentration of 10 µg/ml and cOmplete^TM^ EDTA free protease inhibitors, 1 tab 50 ml. Cells were lysed by sonication and centrifuge at 22,000 rpm for 50 mins at 4°C. Clarified lysates were incubated with 5ml HisTrap^TM^ HP (Cytiva #17-5248-02) and washed with 40 CV lysis buffer, then 35 CV of a high salt buffer containing 20 mM Tris (pH 8.0 at 4°C), 1 M NaCl, 35 mM imidazole, 10 mM MgCl, 50 mM KCl, 2 mM ATP and 5 mM βME, then 20 CV of lysis buffer before elution with lysis buffer containing 500 mM imidazole.

6His-GFP-Mis18_FL_ and 6His-GFP-Mis18_121-194_ were lysed in lysis buffer containing 20 mM Tris (pH 8.0 at 4°C), 100 mM NaCl, 35 mM imidazole and 5 mM βME and supplemented with DNase to a concentration of 10 µg/ml and cOmplete^TM^ EDTA free protease inhibitors, 1 tab 50 ml. Cells were lysed by sonication and centrifuged at 22,000 rpm for 50 mins at 4°C. Clarified lysates were incubated with 5ml HisTrap^TM^ HP and washed with 40 CV of lysis buffer, then 35 CV of high salt buffer containing 20 mM Tris (pH 8.0 at 4°C), 1 M NaCl, 35 mM Imidazole, 10 mM MgCl, 50 mM KCl, 2 mM ATP and 5 mM βME, then 20 CV of lysis buffer before elution with a lysis buffer containing 500 mM imidazole. The proteins were concentrated and SEC was preformed on an ÄKTA system using a S200 increase 10/300 column (Cytiva) equilibrated with 20 mM Tris (pH 8.0 at 4°C), 150 mM NaCl, and 2 mM DTT.

### Protein binding assays

Amylose resin (NEB cat #E8021S) was washed with binding buffer (20 mM HEPES pH 7.0, 150 mM NaCl, 1 mM TCEP, 0.01% Tween-20). 6His-MBP or 6His-MBP-Sad1_2-167_ were mixed with candidate binding proteins and resin with each protein at 5 μM. “Total” samples were taken, and reactions were incubated at 4°C for 1 hr with gentle rotation. Resin was then pelleted and washed in 4 exchanges of 500 μL binding buffer. All supernatant was removed, sample buffer was added, and samples were boiled at 95°C for 3 minutes to yield the “Pulldown” sample. Equivalent proportions of the total reaction from “Total” and “Pulldown” samples were loaded on SDS-PAGE gels (NuPage, cat #NP0322BOX), which were stained with Imperial Protein Stain (Coomassie R-250, ThermoFisher, cat #24615).

### qChIP

For endogenous centromere CENP-A ChIP assays, replicate cultures were grown in YES media (3+ cultures of the same genotype). *sad1-1*-containing strain starter cultures were grown at 25°C to log phase and used to inoculate sample cultures at 25°C or at 36°C for 8 hrs before fixation. For Sad1-TEV140-3V5 ChIP, starter replicate cultures were grown in YES, then washed in media without thiamine, and used to inoculate PMG complete media with or without 15 μM thiamine and cultured for 10 hrs at 32°C. Efficient Sad1-TEV140-3V5 cleavage was assayed by western. For CENP-A establishment assays, fresh pHcc2 transformants were cultured in PMG -adenine-uracil media to log phase. Before fixation, a sample of cells was plated on selective media, then replica -plated to YES low adenine media (10 μg/mL adenine) to test for red/white colony sectoring. Pure white colonies on low adenine indicates plasmid integration, so samples harvested from strains with a high proportion (>10%) of pure white colonies were omitted from further processing. In all cases, cells were fixed at approximately OD_600_ = 1.0 by addition of formaldehyde (Merck, CAS# 50-00-0) to 1% followed by 15 min incubation at RT. Crosslinking was quenched with 125 mM glycine for 5 min. Cells were pelleted, washed in phosphate buffered saline (PBS), and snap frozen in liquid nitrogen.

### Immunoprecipitation

For coIP and MS experiments, cells were grown to log phase (approximate OD_600_ = 2.0) in YES or 4xYES, pelleted, resuspended in SPB lysis buffer with protease inhibtors and phosphatase inhibitors, and flash frozen as pellets. SPB lysis buffer was 25 mM HEPES, 150 mM KCl, 2 mM MgCl2, 0.5% Triton-X 100, 10% glycerol, and was supplemented with 1 mM DTT. Protease inhibitors were 1 mM PMSF and a cocktail (Sigma, cat #P-8215) used at 1:100 dilution. Phosphatase inhibitors were 10 ng/mL microcystin (Enzo cat #ALX350-012-C100), 1 mM sodium pyrophosphate, 100 μM sodium orthovanadate, 5 mM sodium fluoride, and 2 mM β-glycerophosphate. For Fig. 1 and S1 experiments, lysis was performed with a Retsch MM400 ball mill with 3 rounds of milling at 30 cycles/second for 2 min/cycle with 2 min in liquid nitrogen between rounds. For other MS and IP experiments, cryo-lysis was performed with a freezer mill (Spex 6875) with 8-10 rounds of 2” at 10 cycles /second. Lysate was gently sonicated, treated with 50 u/mL benzonase (EMD Millipore, Cas #9025-65-4) at 4°C for 1 hr, then pelleted at 4,700 g for 10 min at 4°. α-V5 or α-Flag M2 antibodies were conjugated to Protein G Dynabeads (ThermoFisher cat #10009D) by crosslinking with dimethyl pimelimidate (DMP, ThermoFisher cat #21666). Beads were incubated with lysate supernatant at 4° for 2 hrs, then washed with 5 exchanges of SPB lysis buffer, including 1 mM DTT, 0.2 mM PMSF, 1:500 protease inhibitors, and phosphatase inhibitors in the first three washes. For coIP, beads were eluted in Laemmli sample buffer by heating to 95°C for 3 minutes, then transferring supernatant to a new tube and treating with reducing 50 mM DTT. For MS, beads were eluted in 2 sequential rounds with 0.1% RapiGest SF (Waters cat #186001860) in 50mM Tris-HCl pH 8 by incubating at 50°C for 10 minutes with mixing. Eluate from the second round was pooled with that of the first round for analysis and further processing. Samples were analyzed by SDS-PAGE electrophoresis and silver staining (ThermoFisher cat #LC6070) or western blotting.

### Mass spectrometry

IP eluate was reduced with 25 mM DTT at 80°C for 1 min, then denatured by addition of urea to 8M. Sample was applied to a Vivacon 30k MWCO spin filter (Sartorius cat #VN01H21) and centrifuged at 12.5k g for 15-20 minutes. Protein retained on the column was then alkylated with 100 μL of 50 μM iodoacetamide (IAA) in buffer A (8 M urea, 100 mM Tris pH 8.0) in the dark at RT for 20 min. The column was then centrifuged as before, and washed with 100 μL buffer A, then with 2 x 100 μL volumes of 50 mM ammonium bicarbonate (AmBic). 3 μg/μL trypsin (Pierce #90057) in 0.5 mM AmBic was applied to the column, which was capped and incubated at 37° overnight. Digested peptides were then spun through the filter, acidified with trifluoroacetic acid (TFA) to pH <= 2, loaded onto manually-prepared and equilibrated C_18_ reverse-phase resin stage tips (Sigma #66883-U)^79^, washed with 100 uL 0.1% TFA, and stored at -20°C prior to MS analysis.

Peptides were eluted in 40 μL of 80% acetonitrile in 0.1% TFA and concentrated down to 2 μL by vacuum centrifugation (Concentrator 5301, Eppendorf, UK). The peptide sample was then prepared for LC-MS/MS analysis by diluting it to 5 μL by 0.1% TFA. LC-MS-analyses were performed on an Orbitrap Fusion™ Lumos™ Tribrid™ Mass Spectrometer (Thermo Scientific, UK) coupled on-line to an Ultimate 3000 RSLCnano Systems (Dionex, Thermo Fisher Scientific, UK). In both cases, peptides were separated on a 50 cm EASY-Spray column (Thermo Scientific, UK), which was assembled on an EASY-Spray source (Thermo Scientific, UK) and operated at 50°C. Mobile phase A consisted of 0.1% formic acid in LC-MS grade water and mobile phase B consisted of 80% acetonitrile and 0.1% formic acid. Peptides were loaded onto the column at a flow rate of 0.3 μL min-1 and eluted at a flow rate of 0.25 μL min-1 according to the following gradient: 2 to 40% mobile phase B in 150 min and then to 95% in 11 min. Mobile phase B was retained at 95% for 5 min and returned back to 2% a minute after until the end of the run (190 min). FTMS spectra were recorded at 120,000 resolution (scan range 350-1500 m/z) with an ion target of 7.0×105. MS2 was performed in the ion trap with ion target of 1.0×104 and HCD fragmentation^80^ with normalized collision energy of 27. The isolation window in the quadrupole was 1.4 Thomson. Only ions with charge between 2 and 7 were selected for MS2. The MaxQuant software platform^81^ version 1.6.1.0 was used to process the raw files and search was conducted against *Schizosaccaromyces pombe* complete/reference proteome set of PomBase database (released in July, 2016), using the Andromeda search engine^82^. For the first search, peptide tolerance was set to 20 ppm while for the main search peptide tolerance was set to 4.5 pm. Isotope mass tolerance was 2 ppm and maximum charge to 7. Digestion mode was set to specific with trypsin allowing maximum of two missed cleavages. Carbamidomethylation of cysteine was set as fixed modification. Oxidation of methionine was set as variable modification. Label-free quantitation analysis was performed by employing the MaxLFQ algorithm as described^83^. Absolute protein quantification was performed as described^83^. Peptide and protein identifications were filtered to 1% FDR.

Perseus^85^ version 1.6.2.1 was used to analyze output of the MaxQuant searches. Protein identifications based on reverse peptide, potential contaminant, and “only-site” IDs (only modified peptides) were filtered out, as were IDs with less than 2 valid peptide IDs. Intensity values were Log_2_ transformed to facilitate statistical analysis. Replicate data was grouped, and proteins with less than 2 valid values in at least one group were filtered out. Imputation replaced missing values from a normal distribution with settings width = 0.3 and downshift = 1.8. Volcano plots were generated using 2-sample Student’s t-tests with a p-value of 0.05 (alpha = 0.05). Horiontal cutoffs at y = 1.3 on the plots represent statistical significance by this test. Vertical lines were drawn at Log_2_(+/-2) enrichment as a guide for meaningful abundance differences.

### qChIP

3 or more biological replicates of approximately 2.5x10^8^ cells of were prepared of each ChIP sample. Cells were lysed by bead beating in 350 μL ChIP lysis buffer (50 mM HEPES-KOH, pH 7.5, 140 mM NaCl, 1 mM EDTA, 1% (v/v) Triton X-100, 0.1% (w/v) sodium deoxycholate), then sonciated to solubilize chromatin (20 cycles of 30 sec on + 30 sec off in a 4°C water bath Diagenode Bioruptor on “high”). Cellular debris was removed by centrifugation at 13.2k rpm for 10 min at 4°. For α-Cnp1 ChIP, lysate was pre-cleared by incubating with Protein G agarose resin (Roche, CAS #64-17-5) at 4° for 1 hr. Resin was then removed, input sample was taken, and the remaining supernatant was incubated with Protein G resin plus α-Cnp1 (in house serum, used at 4 uL and 25 uL resin per ChIP). For α-V5 and α-GFP ChIP, the preclearing step was omitted and 25 uL of Protein G Dynabeads were used with 3 μL of α-V5 or 2 μL of α-GFP (ThermoFisher, cat #A11122). IP’s were performed overnight at 4°C. Resin or beads were washed with ChIP Lysis Buffer, then the same buffer with 0.5M NaCl, then ChIP Wash Buffer (10 mM Tris-HCl, pH8, 0.25 M LiCl, 0.5% NP-40, 0.5% (w/v) sodium deoxycholate, 1 mM EDTA), then TE (10 mM Tris-HCl, pH8, 1 mM EDTA), and then processed in tandem with input samples. All samples were incubated with a 10% slurry of Chelex 100 (BioRad cat #1421253) at 100°C for 12 min to reverse crosslinks, then cooled to room temperature and incubated at 55°C with 25 μg of Proteinase K for 30 min. Samples were heated to 100°C for 10 min to inactivate Proteinase K. Resin was pelleted and supernatant was collected for ChIP reactions. qPCR was performed in tripilcate for each sample using primers specific for *actin* or centromeric sequence (see Supplemental Table S7) using LightCycler 480 SYBR Green I Master mix (Roche, 04887352001) on a Roche LightCycler 480 Instrument and LightCycler 480 software v. 1.5.1.62. Analysis is described below.

### Immunofluorescence

Cells were fixed in 3.7% formaldehyde for 15 min at RT, then washed 2x in PEM (100 mM PIPES, pH 7.0, 1 mM EDTA, 1 mM MgCl_2_), 1x in PEMS (PEM + 1.2 M sorbitol), and stored in PEM + 0.1% sodium azide. Fixed cells were treated with 1 mg/mL zymolyase T100 (MP Biomedicals cat #08320932) at 37°C for 90 min, washed in PEMS, then permeabilized in PEMS + 1% Triton-X100 for 1 min. Next, cells were washed in PEMS, then 2x in PEM, then blocked for 30 min in PEMBAL (PEM + 1% Bovine Serum Albumin (BSA), 100 mM lysine hydrochloride, 0.1 sodium azide). Cells were incubated with primary mouse α-V5 and sheep α-Cdc11 at 1:1,000 in PEMBAL at 4°C overnight. 3 x 30 minute washes in PEMBAL were followed by secondary donkey α-mouse Alexa-488 (ThermoFisher RRID: AB_2534082) and donkey α-sheep Alexa-594 (ThermoFisher RRID: AB_2534083) incubation, both at 1:1,000 (2 μg/mL) in PEM, at 4°C for 4 hrs in the dark. Folllowing a PEMBAL wash, cells were stained with 2 μg/mL 4’,6-diamidino-2-phenylindole (DAPI) in PEM at RT for 5 min. Finally, cells were washed once and stored in PEM + 0.1% azide at 4°C.

### Microscopy and image analysis

Fixation was performed by adding formaldehyde to 3.7% and incubating at room temperature for 15 minutes. Cells were washed twice in PEM, then 1x in PEMS. Cells were then stored in PEM + 0.01% sodium azide at 4°C. For imaging, fixed cells were mounted on poly-lysine-coated glass slides (Epredia cat #J2800AMNZ) with Vectashield mounting medium (Vector Laboratories cat #H-1200) containing 1.5 μg/mL 4’,6-diamidino-2-phenylindole (DAPI). For IF samples, mounting media without DAPI was used (Vector Laboratories cat #H-1000).

Images were acquired on a Nikon Eclipse Ti2 inverted microscope with a Lumencore Spectra X lightsource (Beaverton, OR, USA) and a Photometrics Prime 95B camera (Teledyne Photometrics, Birmingham, UK) controlled with Nikon Elements v. 5.2. The stage was controlled with a MadCity Nanodrive (Mad City Labs, Madison, WI, USA). Images were acquired using a 100x 1.49 NA CFI Plan Apochromat TIRF objective. Semrock filter sets were used (excitation 378 nm/emission 460 nm; excitation 488 nm/emission 535 nm; and excitation 578 nm/emission 630 nm). Z-stacks of each field were taken at 0.2 uM steps for 11 slices.

To quantify Csi1-GFP and Lem2-GFP intensity, images were max projected in FIJI. Circular 7x7 pixel regions encompassing individual Sid4-mCherry foci were designated as regions of interest (ROIs), and intensity measurements were made in both red and green channels. A background (cell free region) intensity measurement of the same dimensions was subtracted from ROI intensity measurements in both channels. The adjusted green signal channel intensity was divided by the adjusted red signal channel intensisty for each ROI to yield a normalized GFP intensity measurment.

### Quantification and Statistical Analysis

qPCR results were quantified with LightCycler 480 software v. 1.5.1.62 and analyzed with Graphpad Prism v. 9.5 (RRID: SCR_002798). Ct values of triplicate PCRs were averaged and percent IP was calculated as the ratio of the “IP” sample Ct value to the “Input” Ct value. Where indicated, percent IP of the target locus was normalized to percent IP of the actin locus. Comparisons between biological replicates were performed using Welch’s t-test and bar graphs show means, standard deviations, and individual data points.

For microscopy experiments, the number of cells analyzed (n) is given in the figure for each condition. Data was analyzed by χ^2^ test using Graphpad Prism v 9.5.

---

**Figure S1.**
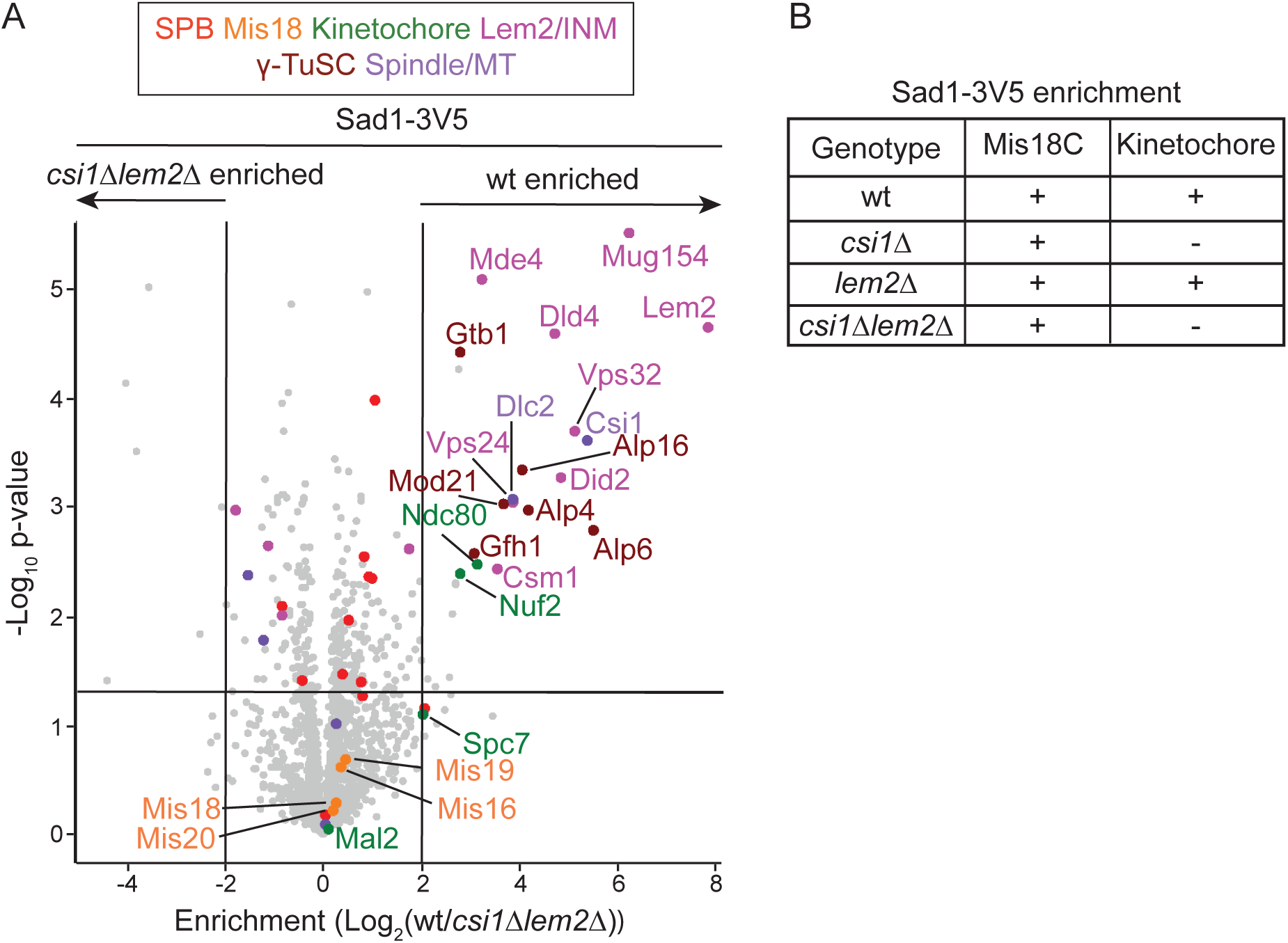
IP/LFQ-MS of Sad1-3V5 in *csi1Δ lem2Δ* shows reduced association of similar proteins as individual *csi1Δ* and *lem2Δ* mutants combined. [Related to Figure 1 and Table S2]. (A) Proteins detected in Sad1-3V5 IPs from wild-type (wt) and *csi1Δlem2Δ* cells were compared by LFQ-MS. Proteins with wt over *csi1Δlem2Δ* enrichments above Log_2_(2) and that were also depleted in either single mutant (Figure 1 D-E) are highlighted and labelled in addition to Mis18 complex and kinetochore proteins. (B) Table indicating whether Mis18 complex or kinetochore proteins are strongly copurified with Sad1-3V5 from cells with the indicated genotypes.

**Figure S2.**
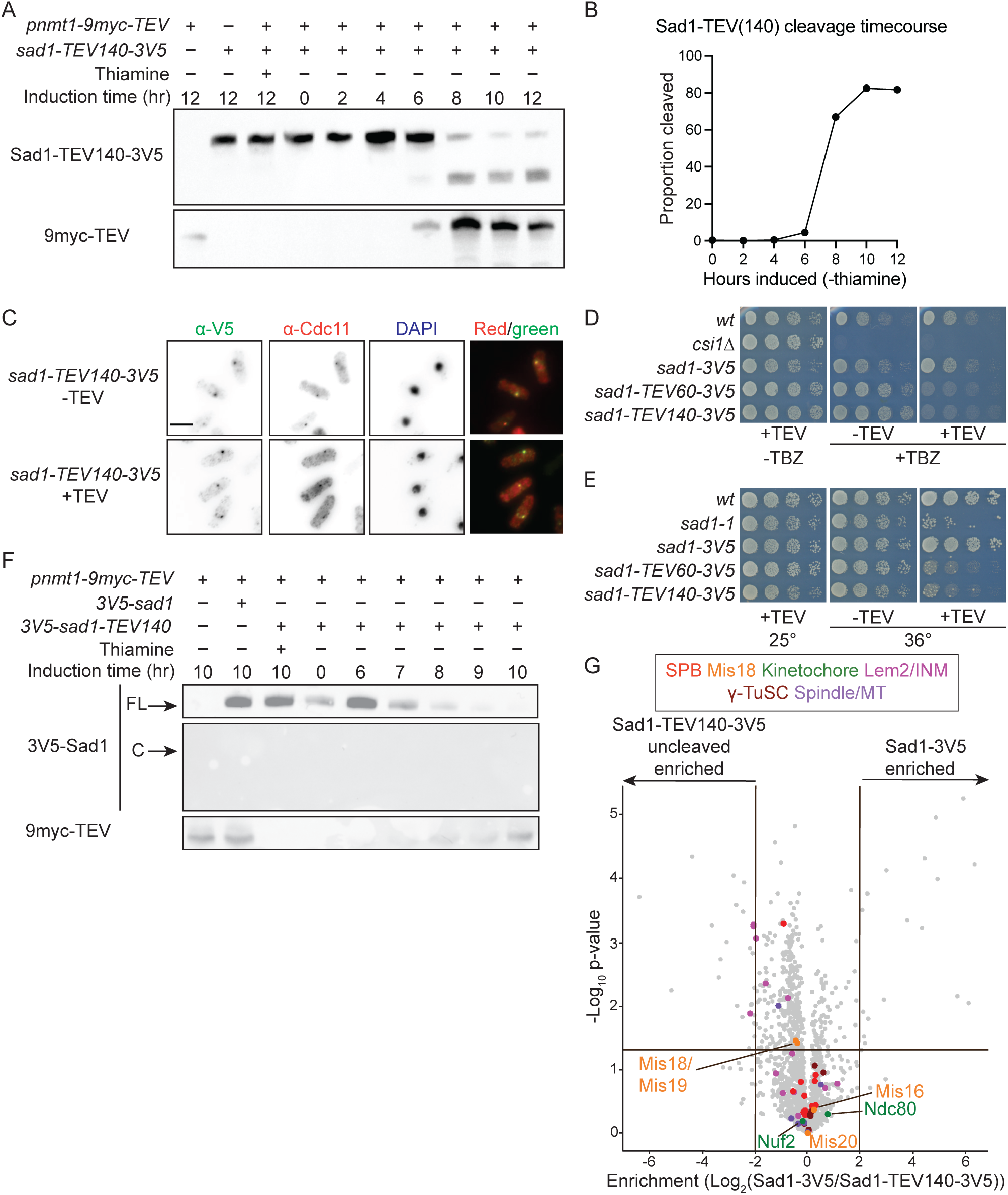
Induced TEV cleavage of Sad1 nucleoplasmic region disrupts specific protein interactions, but not SPB localization of the remaining C-terminal portion of Sad1-TEV140-3V5. [Related to Figure 2 and Table S3]. (A) Detection of Sad1-TEV140-3V5 cleavage (top) and TEV induction (bottom) during a time course by anti-V5 and anti-myc westerns, respectively. Cells from rich media (+thiamine) were washed and placed in +thiamine or -thiamine media at T=0. (B) Quantification of Sad1-TEV140-3V5 cleavage levels from time course in A. Proportion cleaved = cleaved/(cleaved + uncleaved). (C) Sad1-TEV140-3V5 immunolocalization relative to SPBs (anti-Cdc11) with (-) or without (+) TEV cleavage following TEV induction by thiamine removal for 12 hours. Scale bar: 5 μm. (D) Serial dilution growth assays of *sad1-3V5, sad1-TEV60-3V5* and *sad1-TEV140-3V5* cells plated on defined media with (-TEV) or without (+TEV) thiamine that also did (+) or did not (-) include 12.5 μg/mL TBZ at 25 or 36°C. Wild-type (*wt*) and *csi1*Δ included as controls for TBZ sensitivity. (E) Serial dilution growth assays of *sad1-3V5, sad1-TEV60-3V5* and *sad1-TEV140-3V5* cells plated on defined media with (-TEV) or without (+TEV) thiamine incubated at 25 or 36°C. Wild-type (*wt*) and *sad1-1* cells included as controls for temperature sensitivity. (F) Probe for full length (FL) N-terminally tagged 3V5-Sad1-TEV140 (top) with TEV induction (bottom), and the expected N-terminal region (3V5-Sad11-140) cleaved (C) fragment (middle), with anti-V5 and anti-myc westerns respectively. Cells from rich media (+thiamine) were washed and placed in - thiamine media at T=0. (G) Comparison of proteins enriched in Sad1-3V5 and uncleaved Sad1-TEV140-3V5 immunoprecipitates by IP/LFQ-MS.

**Figure S3.**
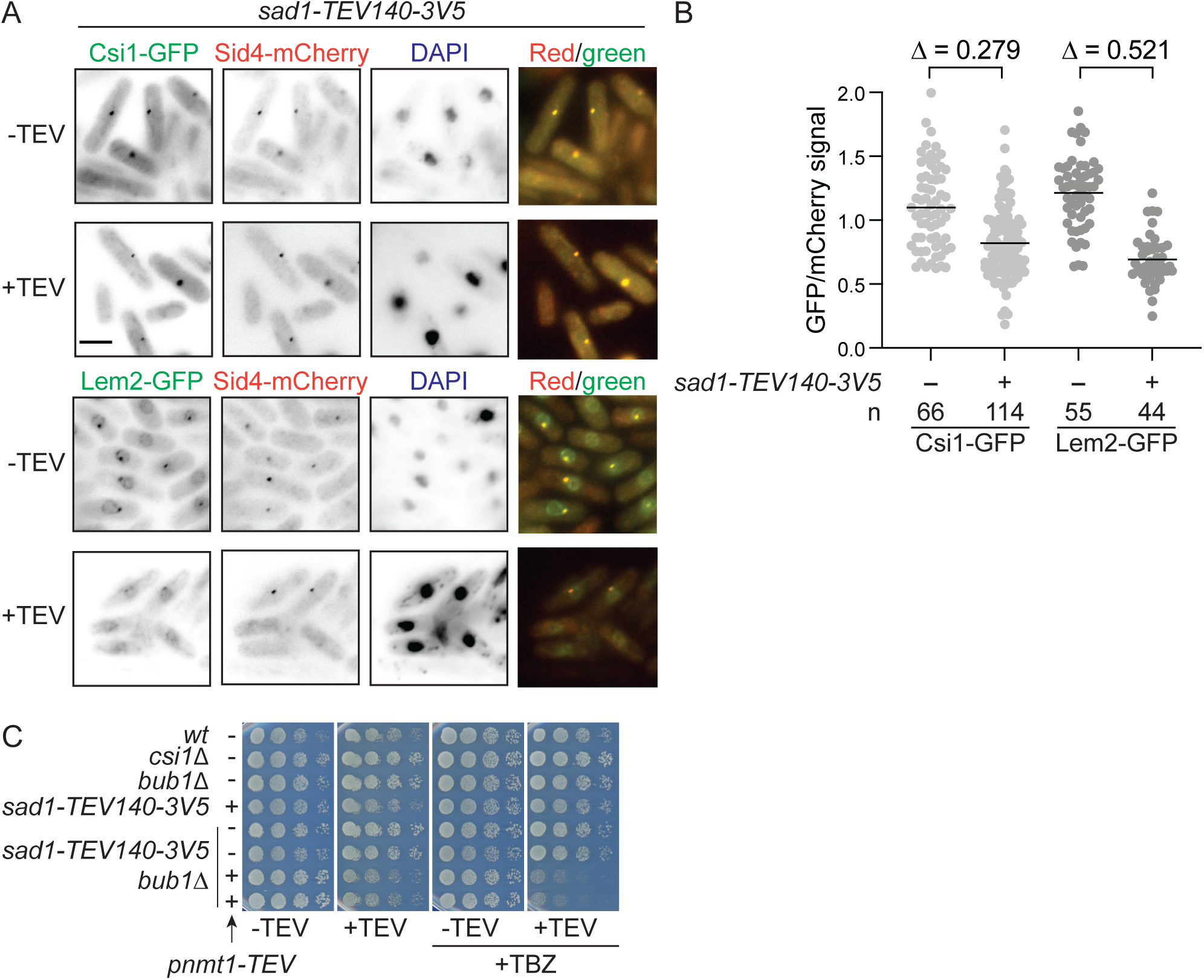
Cells with cleaved Sad1-TEV140-3V5 have aberrant Lem2 localization and are sensitive to Bub1 loss. [Related to Figure 3 and Table S3]. (A) Localization of Csi1-GFP (top) or Lem2-GFP (bottom) relative to SPBs (Sid4-mCherry) in *sad1-TEV140-3V5* cells with (+) or without (-) TEV expression for 16 hrs and stained with DAPI. (B) Intensities of Csi1-GFP (light grey) and Lem2-GFP (dark grey) were measured in strains with or without Sad1-TEV140-3V5 cleavage and normalized to Sid4-mCherry signal. The difference in mean intensities are given as Δ. Scale bar: 5 μm. (C) Serial dilution growth assays on plates with (+) or without (-) TEV expression. *pnmt-TE*V indicates presence (+) or absence (-) of an integrated allele of TEV protease under the *nmt1* promoter. TBZ was used at 7.5 μg/mL.

**Figure S4.**
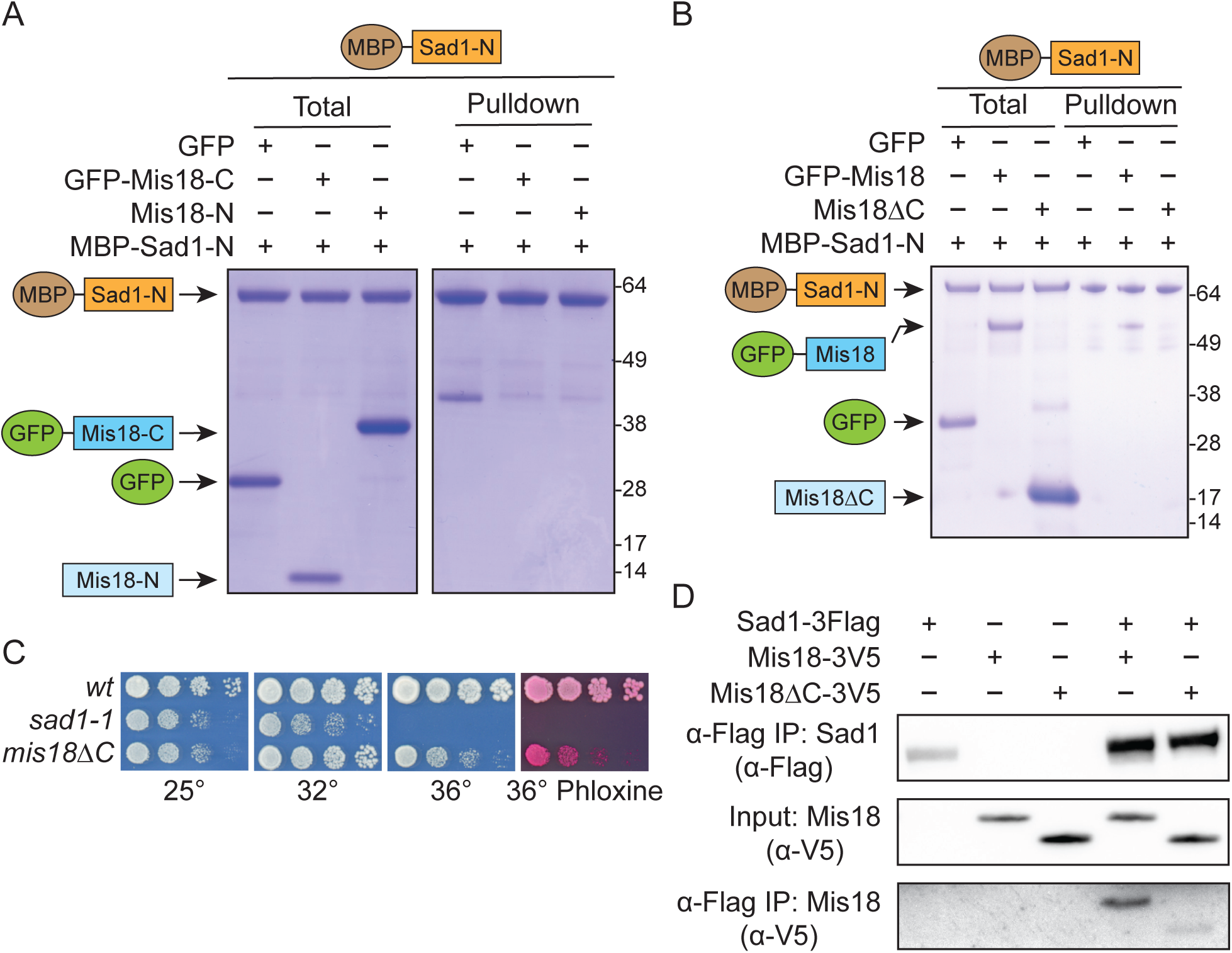
Mis18 truncations fail to bind the Sad1 nucleoplasmic region *in vitro*. [Related to Figure 4]. (A) *In vitro* binding assays for recombinant N-terminal (residues 1-120) and C-terminal (residues 121-194, fused to GFP) portions of Mis18 or GFP (control) to MBP-Sad1-N (residues 2-167, nucleoplasmic region) bound to amylose resin. Size markers: kDa. (B) *In vitro* binding assays for recombinant full length Mis18 fused to GFP, Mis18ΔC (missing C-terminal residues 169-194) or GFP to MBP-Sad1-N (residues 2-167) bound to amylose resin. (C) Serial dilution growth assays of wild-type (*wt*), *sad1-1* and *mis18*Δ*C* cells on YES plates at indicated temperatures and in the presence of phloxine, where red staining indicates inviable cells. (D) Western analyses of Input (extract) and anti-Flag Sad1-3Flag IPs to detect Sad1-3Flag (anti-Flag), Mis18-3V5 or Mis18ΔC-3V5 (anti-V5).

**Figure S5.**
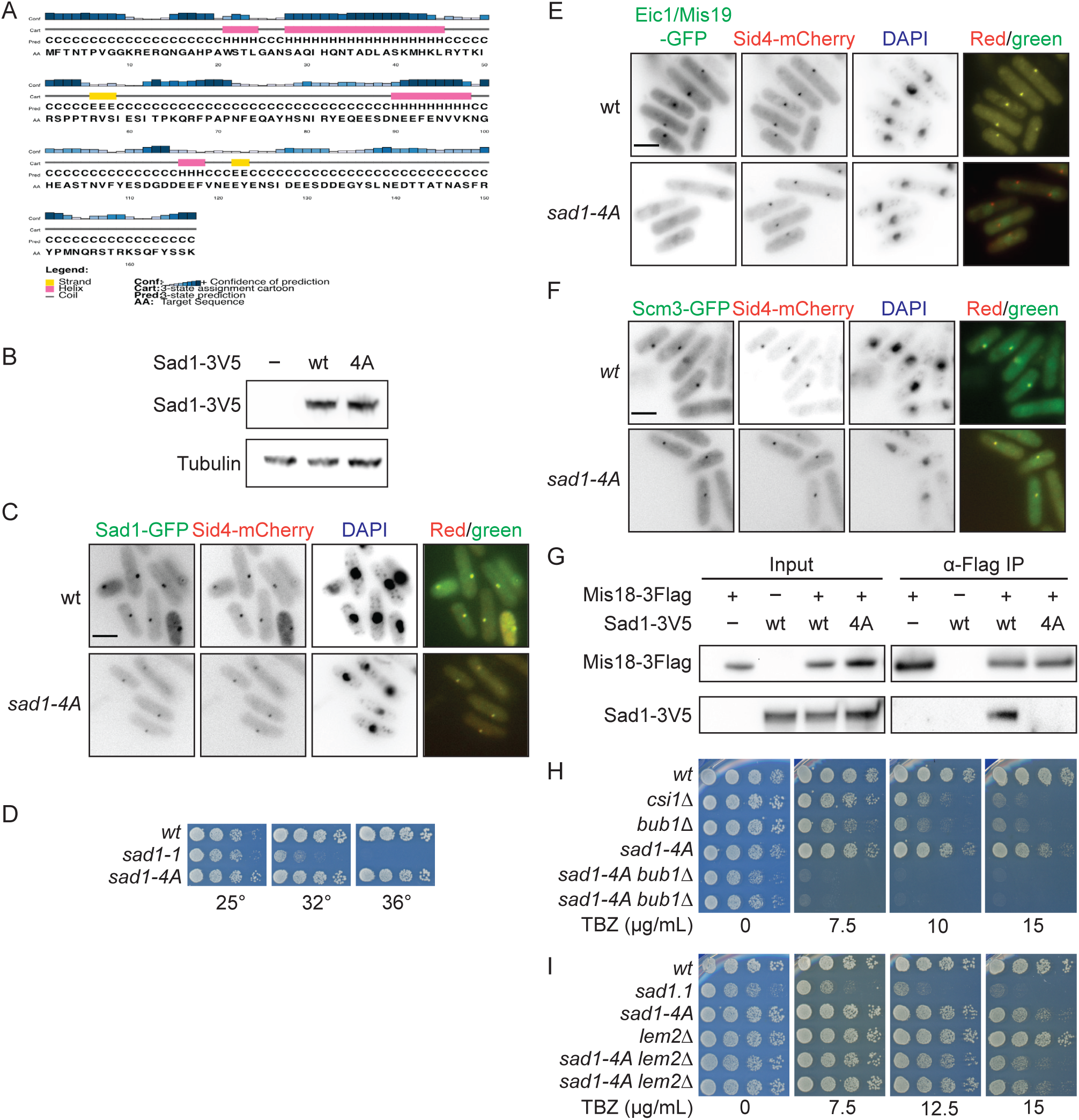
The *sad1-4A* mutant disrupts localization of Mis18C components to SPBs and Mis18-Sad1 association, and exhibits specific genetic interactions. [Related to Figure 5 and Table S4]. (A) PSIRPRED analysis of the Sad11-167 nucleoplasmic region identifies a helical domain covering residues 21-24. (B) Western analysis comparing levels of Sad1-4A-3V5 and Sad1-3V5 protein expression using anti-V5 (α-V5) and anti-tubulin (Tat1). (C) Sad1-GFP and mutant Sad1-4A-GFP protein localization with SPBs (Sid4-mCherry) in cells stained with DAPI. Scale bar: 5 μm. (D) Serial dilution growth assays on YES media at 25, 32, and 36°C. (E) Localization of Mis18C component Eic1/Mis19-GFP and SPB protein Sid4-mCherry in wild-type *sad1^+^*and mutant *sad1-4A* DAPI stained cells. Scale bar: 5 μm. Cells were grown, imaged, and quantified along with those in Figure 5B. (F) Localization of Scm3-GFP and SPB protein Sid4-mCherry in wild-type *sad1^+^* and mutant *sad1-4A* DAPI stained cells. Scale bar: 5 μm. (G) Western analyses of Input (extract) and anti-Flag Mis18-3Flag IPs to detect Sad1-3V5 (wt) or Sad1-4A-3V5 (4A) (anti-V5) and Mis18-3Flag (anti-Flag). (H, I) Serial dilution growth assays of indicated strains on YES plates with or without TBZ added at indicated concentrations.

**Figure S6.**
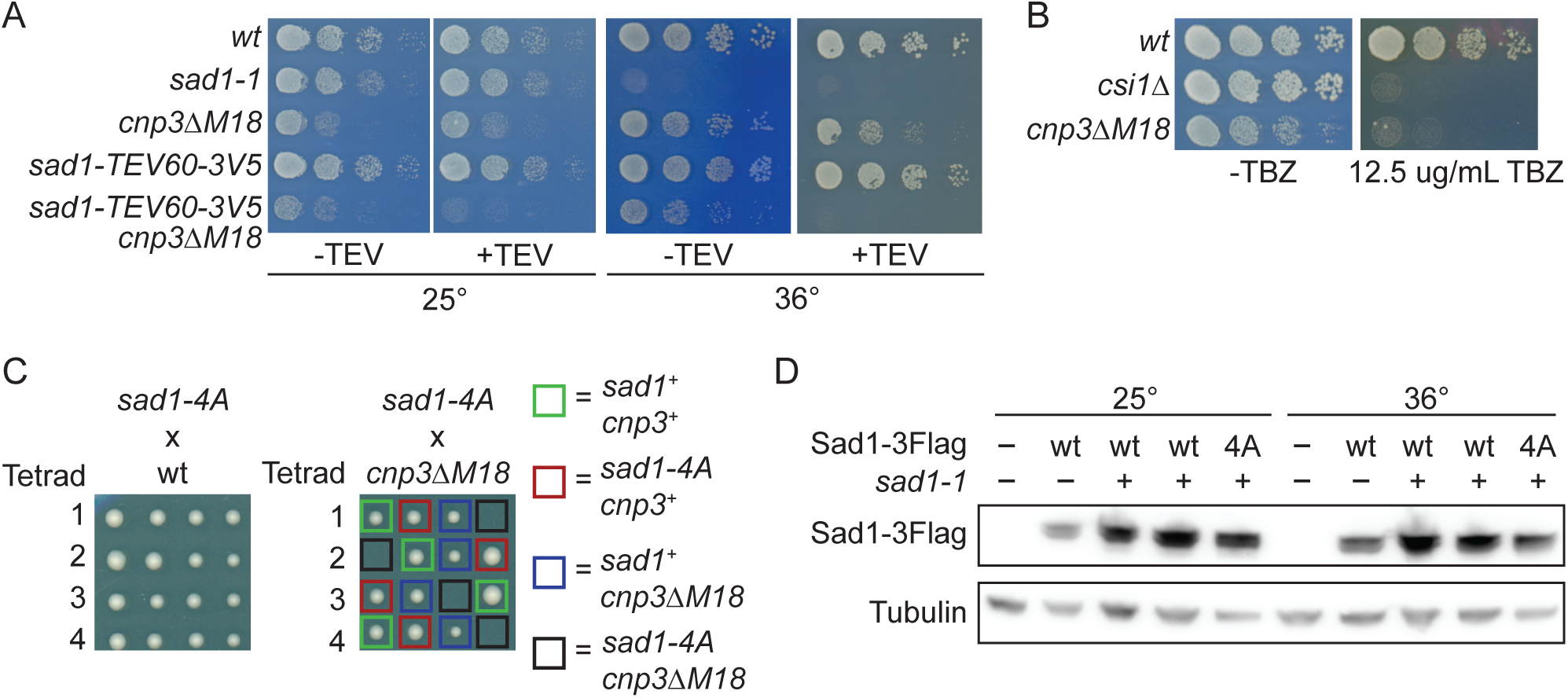
The *cnp3ΔM18* mutant is synthetic lethal with Sad1-TEV60-3V5 cleavage or the *sad1-4A* mutant. [Related to Figure 6]. (A) Serial dilution growth assays of *cnp3ΔM18* mutant cells carrying *sad1-TEV60-3V5* with (+) or without (-) TEV expression on PMG plates at 25 or 36°C. wild-type (*wt*) and *sad1-1* cells provide controls. (B) Serial dilution growth assays of wild-type (wt), *csi1Δ* and *cnp3ΔM18* cells on YES plates with or without TBZ. (C) Growth of dissected spores from four tetrads resulting from crossing *sad1-4A* with wild-type (left) or c*np3ΔM18* (right) cells. PCR genotyping confirmed that viable progeny did not carry both mutations as indicated by the colored boxes. (D) Western (anti-Flag) to compare wild-type Sad1-3Flag (wt) and Sad1-4A-3Flag (4A) protein levels expressed from an ectopic locus (*e*) in cells with (+) or without (-) the *sad1-1* temperature sensitive mutation at the endogenous *sad1* locus. Loading control: anti-tubulin (Tat1).

